# Dynamics of dual specificity phosphatases and their interplay with protein kinases in immune signaling

**DOI:** 10.1101/568576

**Authors:** Yashwanth Subbannayya, Sneha M. Pinto, Korbinian Bösl, T. S. Keshava Prasad, Richard K. Kandasamy

## Abstract

Dual specificity phosphatases (DUSPs) have a well-known role as regulators of the immune response through the modulation of mitogen activated protein kinases (MAPKs). Yet the precise interplay between the various members of the DUSP family with protein kinases is not well understood. Recent multi-omics studies characterizing the transcriptomes and proteomes of immune cells have provided snapshots of molecular mechanisms underlying innate immune response in unprecedented detail. In this study, we focused on deciphering the interplay between members of the DUSP family with protein kinases in immune cells using publicly available omics datasets. Our analysis resulted in the identification of potential DUSP-mediated hub proteins including MAPK7, MAPK8, AURKA, and IGF1R. Furthermore, we analyzed the association of DUSP expression with TLR4 signaling and identified VEGF, FGFR and SCF-KIT pathway modules to be regulated by the activation of TLR4 signaling. Finally, we identified several important kinases including LRRK2, MAPK8, and cyclin-dependent kinases as potential DUSP-mediated hubs in TLR4 signaling. The findings from this study has the potential to aid in the understanding of DUSP signaling in the context of innate immunity. Further, this will promote the development of therapeutic modalities for disorders with aberrant DUSP signaling.

## 1. Introduction

Reversible phosphorylation and dephosphorylation events serve as regulatory switches that control the structure, activity as well as the localization of the proteins in subcellular space thereby influencing vital biological processes [1, 2]. A coordinated interplay between protein kinases (PKs) and protein phosphatases is crucial to regulate these intracellular signaling events as perturbation events in the basal phosphorylation levels of proteins can lead to undesirable consequences including development of diseases such as cancer [3]. Over the years, over 500 PKs have been reported [4], a majority of which are druggable [5]. On the contrary, protein phosphatases although being essential regulators of signaling, have drawn less attention. Among the protein phosphatases, the dual-specificity phosphatase (DUSP) family of phosphatases are the most diverse group with a wide-ranging preference for substrates. A unique feature that distinguishes DUSPs from other protein phosphatases is their ability to dephosphorylate both serine/ threonine and tyrosine residues within the same substrates [6]. Recent studies have suggested that there are about 40 members of the DUSP family and nine subfamilies [7]. These DUSPs have been implicated as critical modulators of several important signaling pathways that are dysregulated in various diseases.

DUSPs were initially referred to as Mitogen-activated protein kinase phosphatases (MKPs) and their role as regulators of MAPK signaling mediated cellular processes in both innate and adaptive immunity have been widely discussed [8-8]. For instance, DUSP1 also known as MKP-1 has been found to be a primary regulator of innate immunity [11] and was identified as an important regulator of T cell activation [12]. Additionally, it was also shown to regulate IL12-mediated Th1 immune response through enhanced expression of IRF1 [13]. Upon LPS treatment, DUSP1-deficient mouse macrophages showed increased expression and activation of p38MAPK leading to increased production of chemokines such as CCL3, CCL4, and CXCL2 thereby increasing the susceptibility to lethal LPS shock [14]. In the same study, DUSP-deficient murine macrophages primed with LPS showed transient increase in JNK activity, elevated levels of pro-inflammatory cytokines and increased p38MAPK activation. Further, DUSP10-deficient mice induced with autoimmune encephalomyelitis showed reduced incidence and severity and prevented LPS-induced vascular damage by regulation of superoxide production in neutrophils [15] indicating its key role in innate and adaptive immune responses. Similarly, other DUSPs such as DUSP2 and DUSP5 have been found to participate in the positive regulation of inflammatory processes [16] and are required for normal T cell development and function [17].

In addition to their potential role in immune regulation, studies have implicated the association of DUSPs namely DUSP1, DUSP4 and DUSP6 in oncogenesis especially in the epithelial-to-mesenchymal transition of breast cancer cells and the maintenance of cancer stem cells [18]. Inhibition of DUSP1 and DUSP6 induces apoptosis of highly aggressive breast cancer cells through the increased activation of MAPK signaling [19]. Furthermore, DUSP1-deficient mice form rapidly growing head and neck tumors causing increased tumor-associated inflammation [20]. In addition to members of MKP subfamily, members of the Protein tyrosine phosphatase type IV subfamily (PTPIV, also known as PRLs) have also been suggested to be potential anti-tumor immunotherapy targets due to their role in carcinogenesis with antibody therapy against PRL proteins inhibiting metastasis in PRL-expressing tumors [21, 22]. Additionally, PTP4A3 (PRL-3) has been reported to trigger tumor angiogenesis through the recruitment of endothelial cells [23]. Owing to their regulatory roles in cancer and immunological disorders, DUSPs have been identified as promising therapeutic targets of these diseases [24].

Although the role of certain members of DUSPs have been well characterized, the mechanism by which other members especially atypical DUSPs modulate immune response is still largely unknown. Furthermore, the interplay between members of DUSP family and PKs and their reciprocal actions is minimally understood. Systems biology and integrative biology offer several approaches to identify molecular mechanisms operating behind biological processes in unprecedented detail. Integrated approaches such as Proteogenomics can provide macro-resolution snapshots to facilitate understanding of intricate molecular mechanisms in cancers [25, 26] and infectious diseases [27, 28]. Applying integrated approaches in the context of immunology can therefore offer unique insights into mechanisms of innate and adaptive immunity. In the past few years, several high-throughput datasets were published on naïve and activated immune and hematopoietic cells [29-29]. In this study, an integrated meta-analysis of high-throughput omics datasets related to innate immunity was carried out to delineate the expression dynamics of DUSPs in hematopoietic cells. Additionally, the signaling crosstalk between DUSPs with the members of the protein kinase families in immune cells was deciphered. Finally, we analyzed the association of DUSP signaling pathways downstream of TLR4 signaling Collectively, this study provides potential DUSP-mediated signaling pathways and hubs thus facilitating better understanding of DUSP signaling in innate immunity.

## 2. Results

### 2.1 DUSP classification into subfamilies and evolutionary conservation

We compiled a list of DUSPs from previously published studies and used it to for further analysis. We also aligned protein sequences for known DUSPs, classified them into subfamilies according to their clustering patterns and validated the classification using domain analysis. The classification of DUSPs based on sequence similarity was performed by multiple sequence alignment analysis of DUSP protein sequences (**Figure 1a**). For the analysis, the list of dual specificity phosphatases and their subfamilies was compiled from a recent study [7]. The results were found to be concordant with the classification system described by Chen *et* al [7]. The classification includes members belonging to CDC14, DSP, DSP14, DSP15, DSP3, DSP6, DSP8, PRL, and Slingshot subfamilies. Protein domain analysis of all DUSP members also validated the sub-classification of DUSPs (**Figure 1b**). Most DUSP subfamily members exhibited similar architectures with a common DSPc domain. However, members of a few subfamilies contained additional domains besides DSPc namely CDC14 subfamily (N-terminal DSP domain), DSP1, DSP6, and DSP8 (Rhodanese-like domain) subfamilies and members of Slingshot subfamily (DEK domain at the carboxy terminus). Next, we aimed to determine the evolutionary conservation of dual specificity phosphatases across eukaryotic species by calculating the number of orthologs for all human DUSPs obtained from Homologene database [39]. Our analysis revealed distribution of DUSPs ranging from a minimum, of six orthologs for DUSP2 to 20 for DUSP12 (**Figure 1c**). The distribution of the entire human proteome was similar with a minimum of one and a maximum of 21 ortholog counts. Most DUSPs were found to be conserved in mammals, and particularly among primates suggesting evolutionary conservation across eukaryotic species.

**Figure 1:**
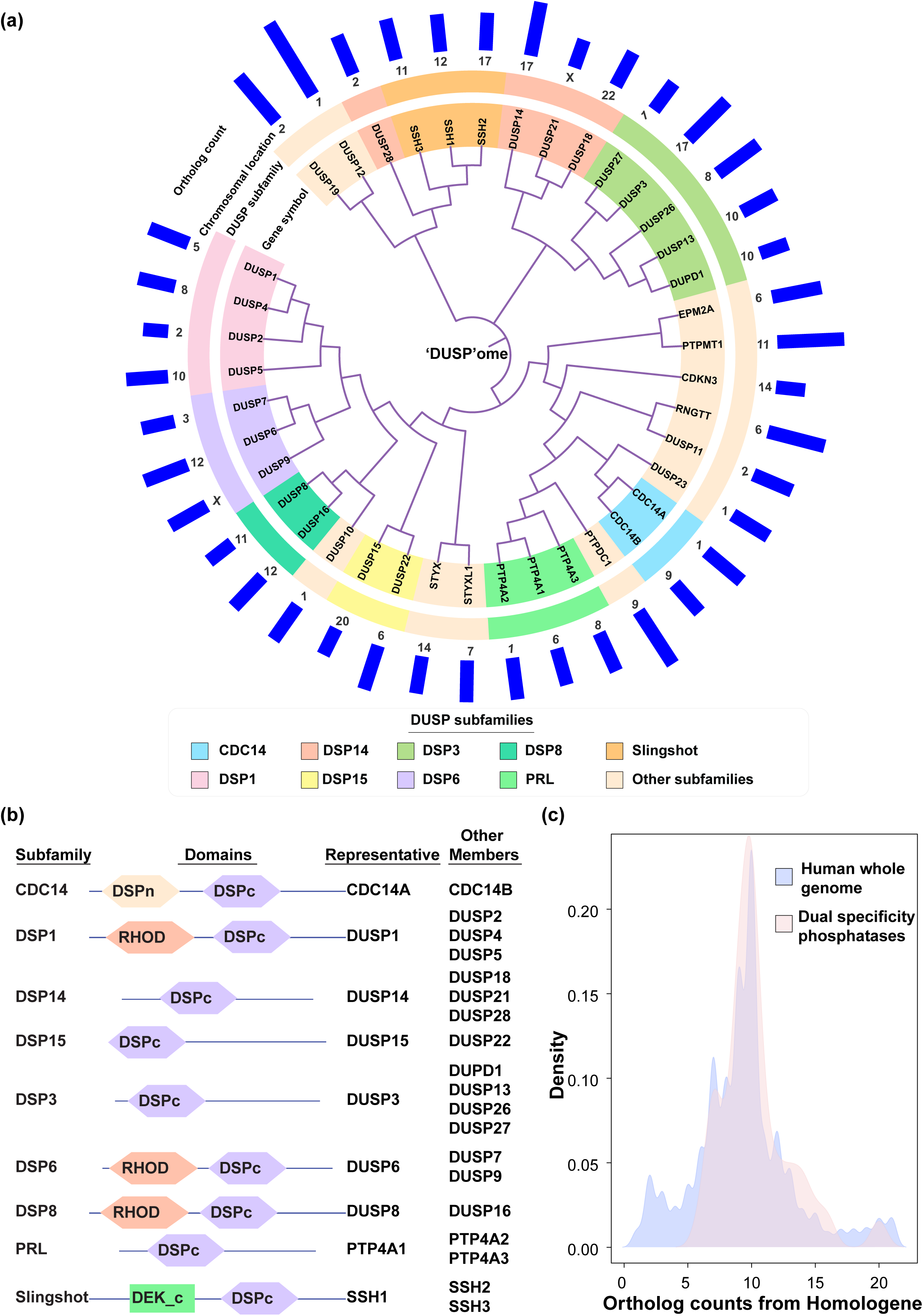
(a) Dendrogram describing the sequence similarity of members of the dual specificity phosphatase (DUSP) family. Protein sequences for various members of the dual specificity family obtained from RefSeq were aligned using Clustal Omega. The DUSPs were classified into subfamilies and their chromosomal location mapped using data and classification system from Chen et al (2017), Science Signaling. The tree shows largely distinct clustering of distinct DUSP subfamilies. **(b) Domain architecture of DUSP subfamilies.** Members of the DUSP family were subjected to domain analysis using SMART domain prediction. **(c) Conservation of dual specificity phosphatases across species.** Ortholog counts were obtained for all human genes from Homologene and the density of ortholog counts for DUSP family members was plotted against the density of ortholog counts for all human genes in the background. The graph largely indicates conservation of DUSPs across various species.

### 2.2 Expression of dual specificity phosphatases and protein kinases in hematopoietic cells, **primary and secondary lymphoid organs**

Earlier reports suggest that the expression of DUSPs are regulated during development in a cell type specific manner or upon cell activation in contrast to their ubiquitous substrate expression [8]. In addition, PKs play important roles in immunity [40, 41] and are widely known to be modulated by DUSPs. In order to determine the extent of expression of DUSPs and PKs across human hematopoietic cells, we analyzed the proteomic data from Rieckemann *et al* [29] as it is currently the largest high-resolution dataset containing expression data pertaining to 28 different hematopoietic cell types analyzed on a single platform (**Supplementary Table S4**, **Supplementary Figure 1**). On an average, 15 DUSPs and 240 PKs were found to be expressed across hematopoietic cells (**Figure 2a**) at the protein level. Among the various cell lineages, T8 TEMRA (terminally differentiated effector memory T cells which express CD45RA, as opposed to TEM cells which are CDC45RA-negative) cells expressed the highest number of DUSPs (19), while erythrocytes did not express any. On the contrary, 264 PKs were found to be expressed in NK CD56^bright^ cells, whereas only 15 were found to be expressed in erythrocytes. NK CD56^bright^ cell types have been previously described to be regulatory in nature influencing innate immunity through cytokine production as opposed to NK CD56^dim^ cells, which have cytotoxic activity [42].

**Figure 2.**
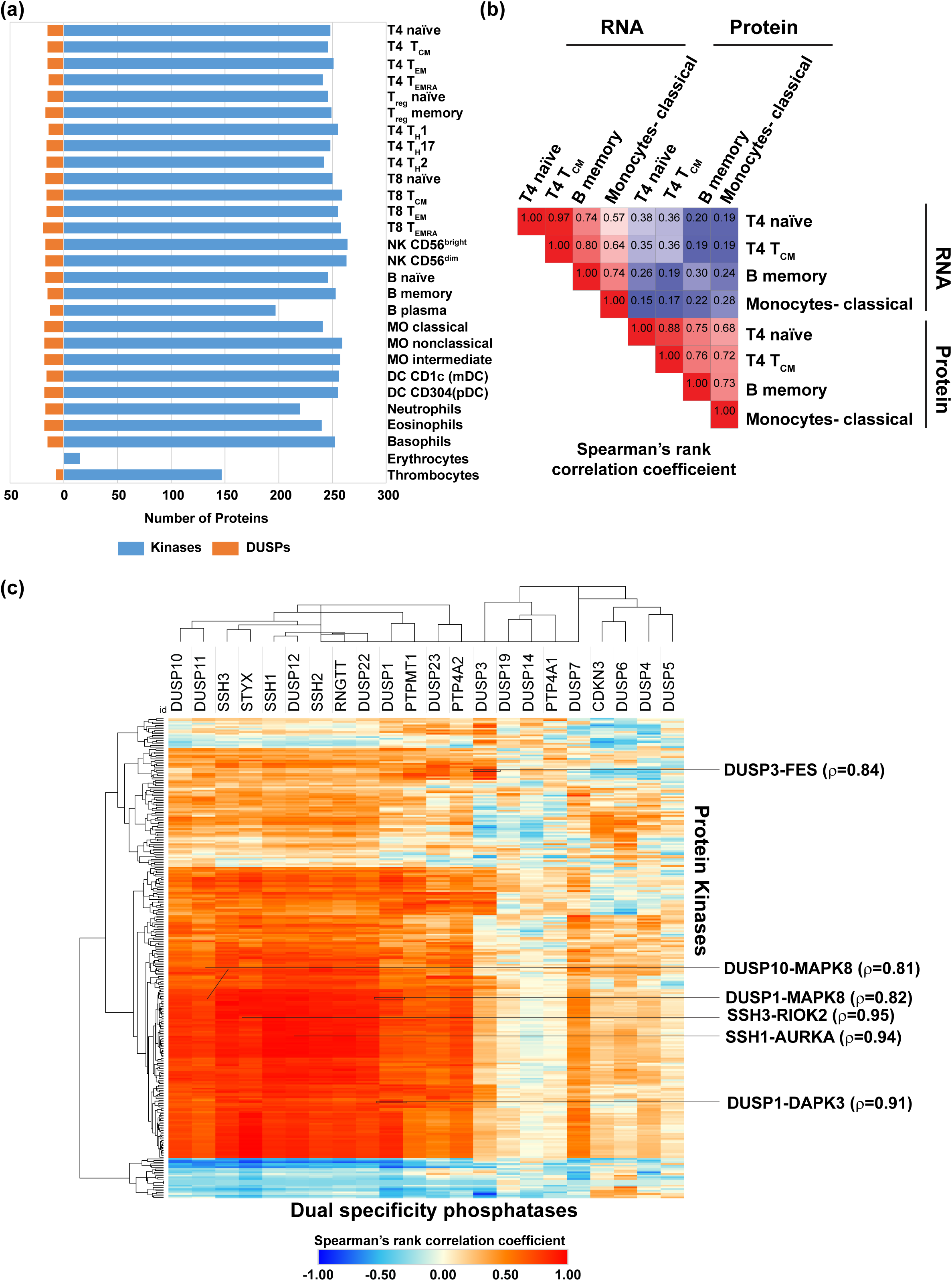
(a) Protein kinases and dual specificity phosphatase expression in hematopoietic cells. Hematopoietic cell expression data for protein kinase and dual specificity phosphatases (DUSPs) were obtained from Rieckmann *et al, Nat Immunol* (2017). All hematopoietic cells except erythrocytes and thrombocytes expressed similar number of kinases and DUSPs. **(b) Correlation of transcriptomic and proteomic data of cells.** Transcriptomic and proteomic data for T4 naïve, T4 TCM, B memory and classical monocytes from Rieckmann *et al, Nat Immunol* (2017) showed poor correlation. **(c) Correlation of protein kinase and DUSP expression patterns in hematopoietic cells**. The correlation was carried out using Spearman’s rank correlation to identify kinase-DUSP pairs that may have reciprocal activities. The kinase-DUSP pairs with high correlation coefficients are shown on the right-hand side.

Among the DUSP family members, 11 DUSPs were found to be expressed in a large majority of the hematopoietic cells analyzed. These include DUSP12, DUSP23, DUSP3, SSH3 and PTP4A2, which were found to be expressed in 27 of 28 hematopoietic cells and DUSP11, DUSP22, PTPMT1, RNGTT, SSH1 and SSH2 found to be expressed in 26 of the 28 cells. 18 members of the DUSP family were not identified in any of the cell types. Furthermore, we did not observe restricted expression of DUSP members to any one cell type. Among the PKs, only 287 were found to be identified in at least one hematopoietic cell type with the rest not being identified in any cell type. While 13 members of the PKs including MAPK1, MAP2K2, ILK, and ROCK2 were found to have ubiquitous expression, four kinases including PRKCD, KALRN, MYLK and PTK2 were found to have restricted expression in thrombocytes. At least five PKs including MKNK2, STK33, CSNK1E, TAOK2 and BLK were found to be restricted to 2 of the 28 cell types. Of these, MKNK2, STK33 and BLK were expressed commonly in B memory cells. STK33 was expressed in B naïve and B memory cells and seemed to be linked with B cell types. We compared proteomics and transcriptomics datasets to see if changes at the transcript level could be equated with changes in protein expression (**Figure 2b**, **Supplementary Table S5**).

We analyzed tissue expression levels of DUSPs and PKs in immune-related tissues. We chose to study expression data pertaining to lymphoid organs that are the major components of the immune system involved in producing B- and T-cells (primary) and are responsible for the coordinating the cell-mediated immune response (secondary) [43]. The primary lymphoid organs consist of the thymus and the bone marrow while the secondary lymphoid organs consist of lymph nodes, spleen, tonsils, and the mucosa-associated lymphoid tissues such as Peyer’s patches [44]. More recently, the appendix has been deemed to be a lymphoid organ capable of carrying out immunological functions [45, 46]. For the analysis, we compiled gene expression data (**Supplementary Table S6**) from multiple projects including FANTOM5 [47], Genotype-Tissue Expression (GTEx) Project [48], The Human Protein Atlas [49, 50], Illumina Body Map [51], NIH Roadmap Epigenomics Mapping Consortium [52] and the ENCODE project [53]. The PK and DUSP expression in these various tissue expression datasets largely correlated **(Figures 3a and 3b).** A majority of the 505 PKs were found to be expressed in at least one lymphoid organ and expression of only a mere 7 kinases were not observed. A significant observation was that all the members of the DUSP family were present in at least one lymphoid organ. 460 of the 505 PKs were found to be expressed in all 7 lymphoid organs while 36 of the 40 DUSPs were expressed in as many lymphoid organs. Two kinases-EPHA8 and PAK5 were found to be restricted to only lymphoid organ-spleen and thymus respectively. DUSP21 was found to be restricted in the bone marrow while DUPD1 was found to be restricted to only the thymus and the tonsil. Similarity matrices indicated a similar distribution of kinases and DUSP expression across primary and secondary lymphoid organs suggesting potential reciprocal pairs (**Figure 3a and 3b**). In conclusion, members of the DUSP family was found to be expressed in every lymphoid tissue suggesting DUSP mediated control of protein kinase activity.

**Figure 3.**
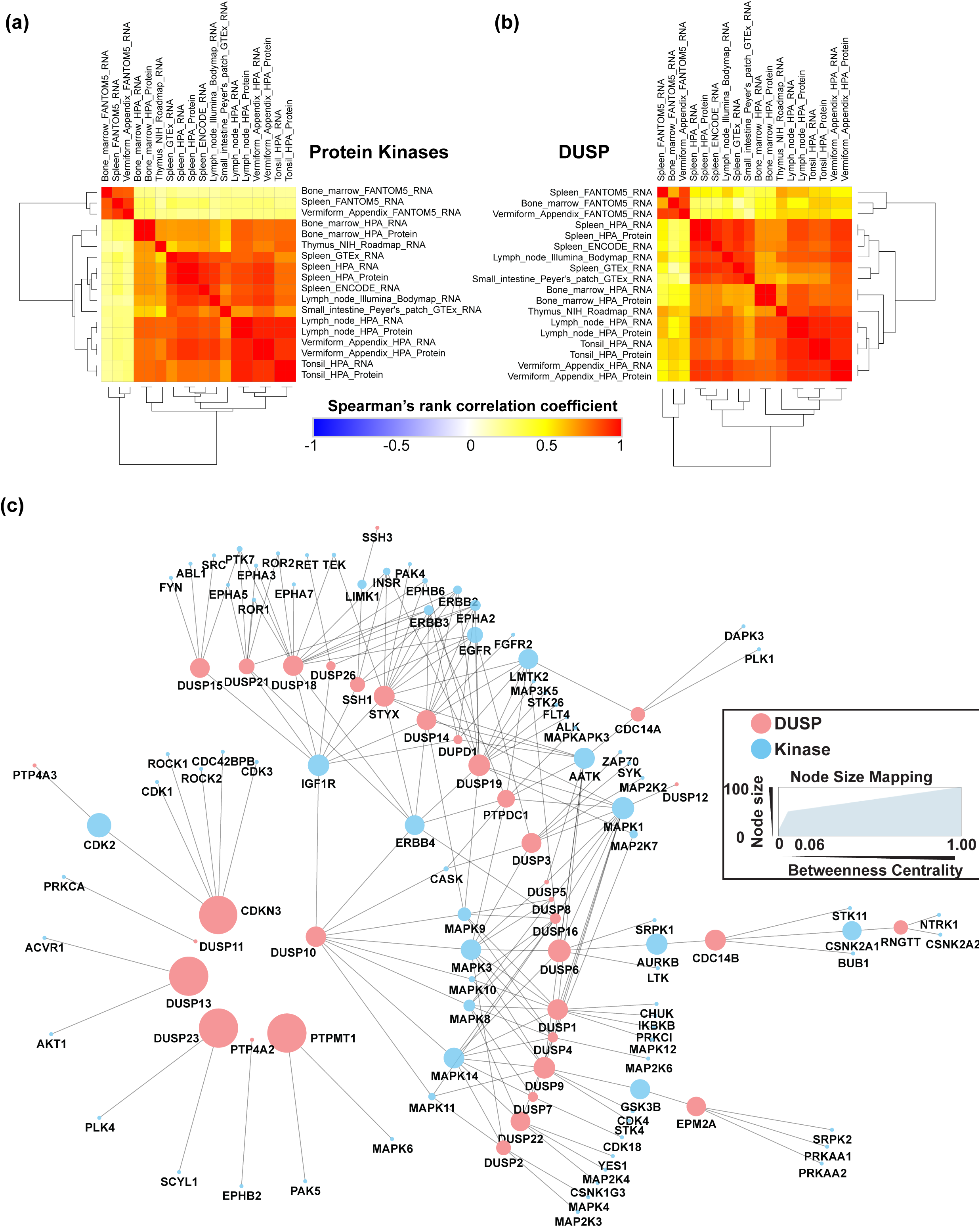
(a) Similarity matrix for protein kinase expression in primary and secondary lymphoid organs. (b) Similarity matrix for DUSP expression in primary and secondary lymphoid organs. The expression data Bone marrow, spleen, thymus, lymph node, tonsil, appendix and Peyer’s patches were obtained from various studies including FANTOM5, HPA, GTEx, ENCODE, Illumina Bodymap and NIH Roadmap project consortia. The PK and DUSP expression in these various tissue expression datasets largely correlated **(c) DUSP-Protein kinase interaction network**. Protein-protein interaction data between DUSP and protein kinases obtained from Compartmentalized Protein-Protein Interaction (comPPI) Database were analyzed in Cytoscape using Network Analyzer to obtain network properties including Betweenness Centrality. Protein kinases with high Betweenness Centrality indicate primary regulatory proteins associated with DUSPs.

### 2.3 Correlation of dual specificity phosphatase activity with protein kinase activity in immune cells

Since it is widely known that DUSPs regulate protein kinases, identifying DUSPs and protein kinases pairs with reciprocal activities could identify regulatory mechanisms that can be potentially exploited to develop strategies for therapeutic interventions in conditions such as systemic inflammatory and autoimmune disorders. To identify potentially reciprocal DUSP-kinase pairs and to determine its probable role in immune cells, we correlated the expression profiles of DUSPs and PKs expressed in hematopoietic cells from Rieckemann *et al* [29] (**Supplementary Table S7**, **Figure 2c)**. We identified 231 DUSP-kinase pairs with similar coexpression patterns in immune cells with a Spearman’s rank correlation coefficient of 0.9 or more indicating high confident reciprocal pairs. Similarly, 701 DUSP-kinase pairs with similar coexpression patterns in immune cells had a Spearman’s rank correlation coefficient of 0.8 or more. Some of the pairs identified include SSH1-AURKA (ρ= 0.94), DUSP1-MAPK7 (ρ= 0.79), DUSP1-MAPK8 (ρ= 0.82), DUSP10-MAPK8 (ρ= 0.81). Among these, the role of DUSP1 and DUSP10 as negative regulators of MAPK8 is well known. The significance of SSH1-AURKA pair in immune cells is currently not well described at this time.

We also used an interactome-based approach to identify co-expressed and co-interacting proteins to identify potential DUSP-kinase regulatory mechanisms. Interactome analysis to identify reciprocal DUSP-protein kinase pairs from baseline protein-protein interaction (PPI) data from the comPPI database resulted in the generation of an interaction network containing 1715 DUSP-specific interactions with 1,276 nodes (**Supplementary Figure 2**). Since the network was too complex to comprehend the interplay between DUSPs and PKs, we separated interactions between DUSPs and kinases and generated an additional kinase and DUSP-specific network. This network contained 195 protein-protein interactions between 35 DUSP members and 82 PKs (**Figure 3c**, **Supplementary Table S8).** Several potential hub proteins that communicate with the dual specificity phosphatase family members were deduced from the interaction network. Among the PKs, MAPK1 and MAPK3 had 12 and 10 interactions (directed edges) respectively with DUSPs. Other PKs with a high number of interactions included MAPK14 (9), IGF1R (9), AATK(9) and ERBB4 (8). Among the DUSPs, DUSP18 and DUSP19 had the most number of directed edges -14 each. Other DUSP family members with several directed edges included STYX (13), DUSP 1 (11), DUSP14 (10), DUSP9 (9) and DUSP 10 (9). It is interesting to note that DUSP19, and STYX are some of the poorly characterized members. The PKs with the highest betweenness centrality included MAPK1, IGF1R, AURKB, and AATK. Combining the directed edge and the betweenness centrality data revealed several kinases belonging to the MAPK family and receptor tyrosine kinases such as MAPK1, MAPK3, IGF1R, AATK, MAPK14, ERBB4, LMTK2, MAPK9 and MAPK8 to be strongly associated with DUSPs.

### 2.4 Expression landscape and signaling dynamics of DUSPs and kinases in activated immune cells

The data provided by Rieckemann *et al* also includes steady-state and activated protein expression profiles of 17 cell types which were analyzed to determine the effect of various activating ligands on the expression of DUSPs and PKs (**Supplementary Table S9**). Our analysis resulted in the identification of 152 events of differential expression of 18 DUSPs across 17 cells types (**Supplementary Figure 3a**) Of these, 57 and 95 were found to be overexpressed and downregulated respectively across the activated cell types. Similarly, we identified 2,311 events of differential expression of 269 PKs across 17 cells types (**Supplementary Figure 3b)**. Of these, 1,058 and 1,253 were found to be overexpressed and downregulated respectively across the activated cell types. Several kinases and DUSPs were found to be differentially expressed in multiple activated cell types. The most overexpressed DUSPs included DUSP12 (9 cell types), DUSP23 (7 cell types), DUSP1, DUSP10 (6 cell types each) and STYX (4 cell types), while the most downregulated DUSPs included SSH3 (13 cell types), PTPMT1 (12 cell types), DUSP3 (10 cell types), SSH2 (9 cell types), DUSP1 (8 cell types) and SSH1 (7 cell types). Similarly, among the kinases the most overexpressed included CDK1 (14 cell types), PLK1 (14 cell types), TGFRB1 (13 cell types), RIOK3 (13 cell types) and RIOK2 (13 cell types) while the most downregulated included ATM (15 cell types), MAST3 (14 cell types), PRKACB (13 cell types), MAP3K5, LMTK2 and SYK (12 cell types each).

Several studies have focused on the genome-wide effects of TLR ligands on hematopoietic cells such as monocytes [54]. In the current study, we aimed to determine the specific roles of DUSPs and PKs in TLR4 signaling. An integrated analysis of RNA and protein expression datasets [29, 33, 55] pertaining to dendritic cells (DCs) and monocytes (MOs) activated by LPS was performed (**Supplementary Table S10).** The data were categorised into murine DCs (mDCs), human DCs (hDCs) and human MOs (hMOs) and a list of molecules differentially expressed in response to LPS was generated (**Supplementary Table S11**). 57 proteins including 4 DUSPs and 53 PKs were found to be overexpressed in mDCs while 80 were downregulated (2 DUSPs, 78 PKs) in response to LPS. In hDCs, 50 proteins (4 DUSPs, 46 PKs) and 154 (10 DUSPs, 144 PKs) proteins were found to were found to be overexpressed and downregulated respectively. In the case of hMOs, 35 (1 DUSP, 34 PKs) were overexpressed and 80 (6 DUSPs, 74 PKs) were downregulated.

DUSPs overexpressed in dendritic cells included Dusp1/DUSP1 (mDCs and hDCs), Dusp14, Dusp16, Ptp4a2 (all in mDCs), DUSP5, DUSP7, and DUSP 10 (all in hDCs). Downregulated DUSPs included Dusp3, Dusp19 (mDCs), DUSP4, DUSP11, DUSP12, DUSP23, PTP4A2, PTPMT1, SSH1, SSH2, SSH3, and STYX (hDCs) (**Figure 4a and Supplementary figure 5a**, **5b and 5c**). In hMOs stimulated with LPS, DUSP 10 was overexpressed while DUSP1, DUSP11, PTP4A2, PTPMT1, RNGTT, and SSH3 were found to be downregulated. DUSP1 (Dusp1) seems to be important in both hDCs and mDCs signaling as it was found to be upregulated in both species in response to LPS and is in in concordance with previously published studies on dendritic cells stimulated with LPS [14]. However, it was found to be downregulated in hMOs. The over expression of DUSP1 in dendritic cells identified from our analysis is in concordance with previously published studies on dendritic cells stimulated with LPS. DUSP10 was found to be upregulated in both hDCs and hMOs while in mDCs, it was not found to be differentially expressed. Among differentially expressed genes in dendritic cells, Dusp14 and Dusp16 were exclusively overexpressed in the dendritic cells from mice while DUSP5, DUSP7, and DUSP10 seemed to be exclusive to humans. Taken together, our analysis suggests probable existence of species-specific differential expression of DUSPs in TLR4 signaling. In a recent paper, human and murine macrophages were found to have varying mechanisms of immunometabolism [56]. Funtional analysis of differentially expressed PKs and DUSPs in murine dendritic cells stimulated with LPS showed enrichment of several processes including MAPK cascade, response to reactive oxygen species, cellular senescence an cell migration pathways among others (**Figure 4b)**.

**Figure 4.**
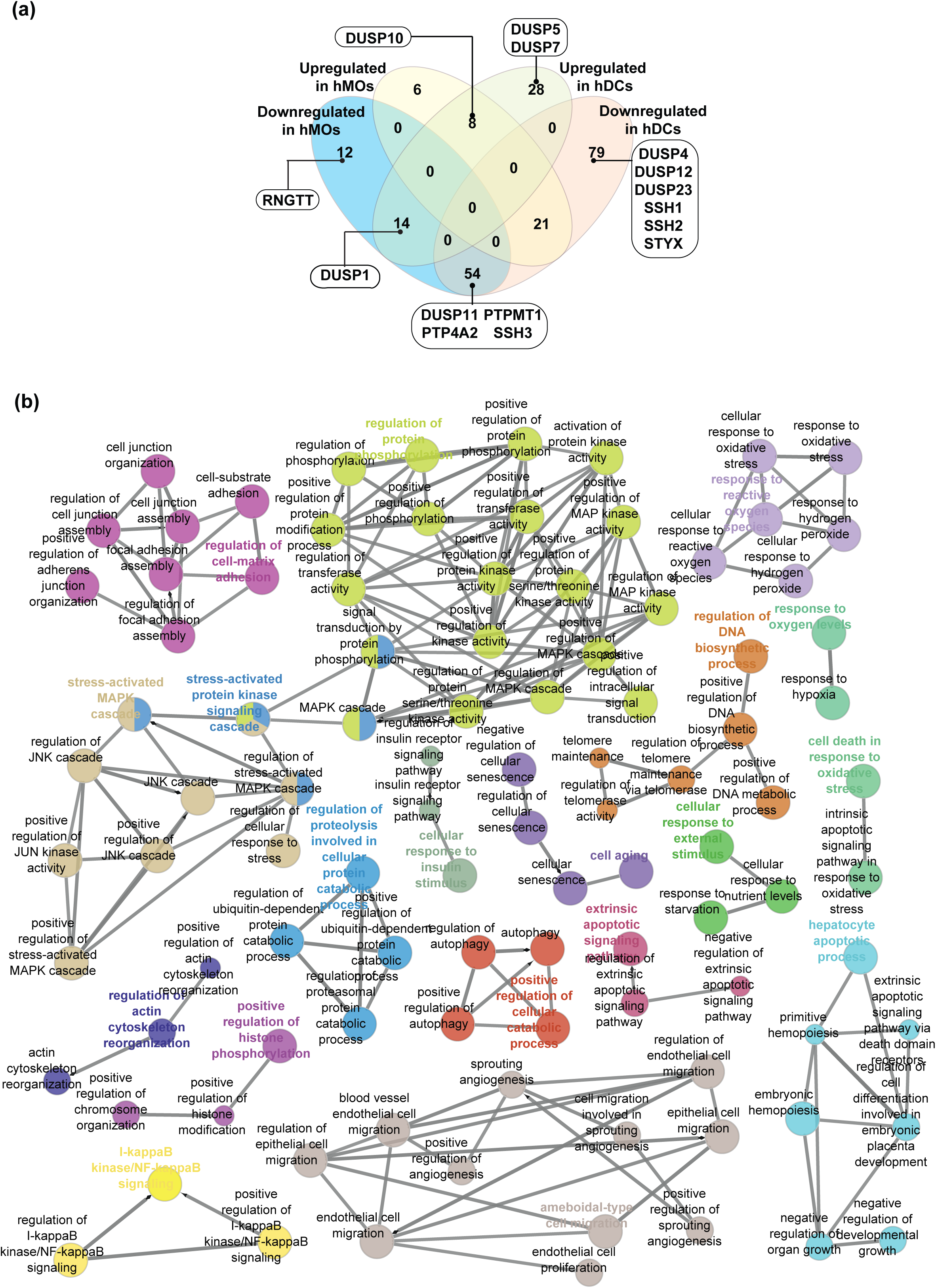
DUSP and kinase dynamics in response to TLR ligands. (a) Venn diagram showing differentially expressed protein/transcripts in human monocytes and dendritic cells stimulated with LPS. Members of the DUSP family are indicated within insets. **(b) Enriched biological processes in murine dendritic cells stimulated with LPS**. Differentially expressed protein kinases and DUSPs in response to LPS were analyzed using ClueGO in Cytoscape. Different colors indicate clusters of similar processes.

**Figure 5.**
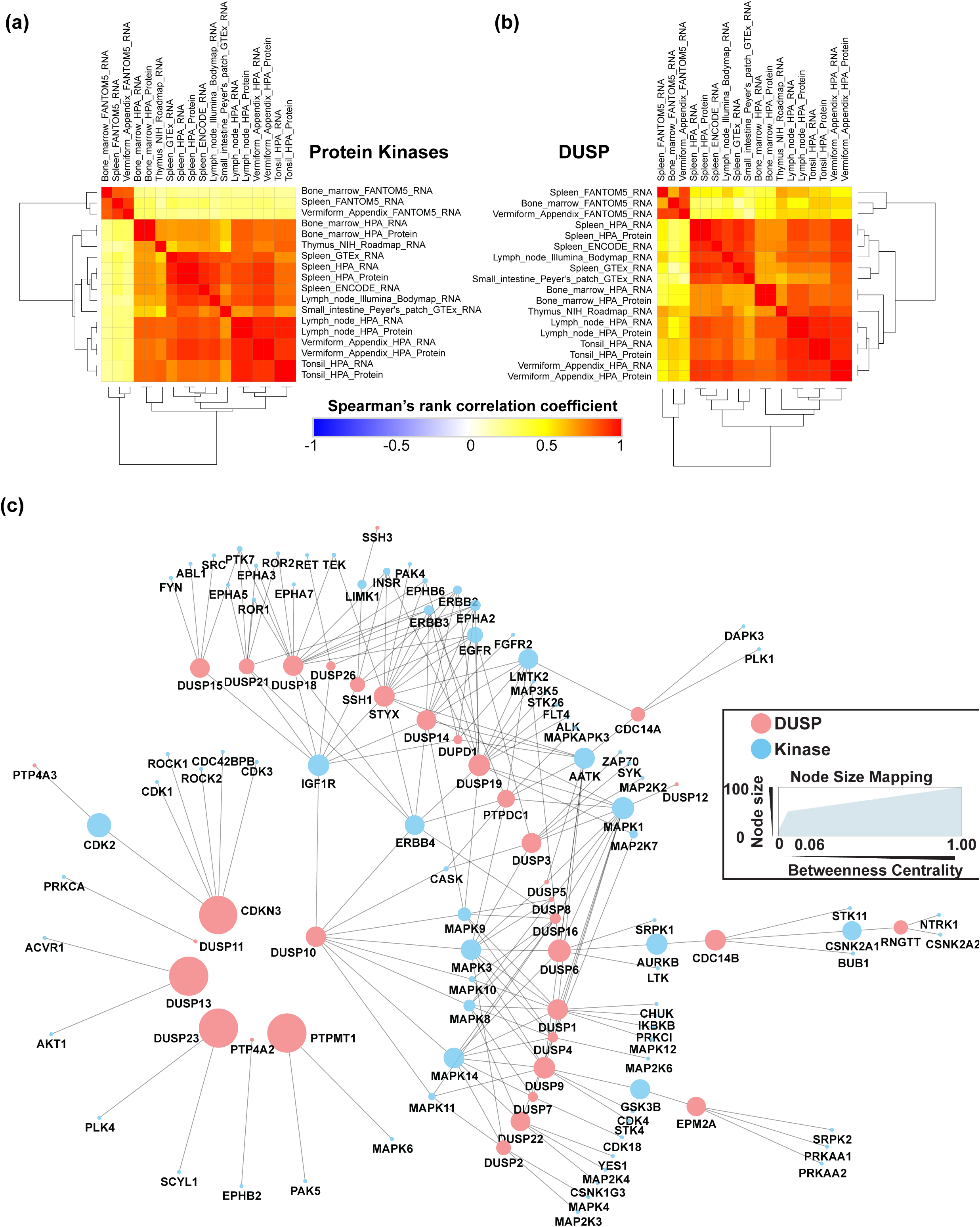
(a) Pathway analysis of DUSPs and kinases differentially expressed in response to LPS. Differentially expressed genes were tested for hypergeometric enrichment of Reactome Pathways. Genesets with less than 10 genes were excluded from the analysis and p-values were adjusted by Benjamini-Hochberg (FDR) correction

We next performed pathway enrichment analysis using pathway data from Reactome database for DUSP and PKs showing differential expression upon LPS stimulation (**Supplementary Table S12**, **Figure 5a**). We found several pathways including TLR signaling, MyD88 cascade, and MAPK signaling pathways to be enriched across LPS-activated MOs and DCs. Oxidative stress-induced senescence pathways were downregulated in mDCs, hDCs, and hMOs. VEGF signaling also seemed to be affected through DUSP and PKs after stimulation with LPS. While induction of TLR signaling by LPS is widely known, there are a few reports on the induction of VEGF signaling by LPS stimulation [57]. FGFR signaling and SCF-KIT signaling pathways were up in hDCs, while C-lectin receptor signaling pathway was downregulated in hMOs upon LPS stimulation. Apoptotic pathways was found to be downregulated in hMOs while being upregulated in hDCs suggesting opposing cell death phenotypes in hMOs and hDCs in response to LPS.

To identify DUSP-kinase pairs with reciprocal activities, correlation analysis between DUSP and kinase expression in DCs and MOs treated with LPS was carried out (**Supplementary Table S13 and Supplementary Figure 5d**). Our analysis resulted in the identification of several important pairs including Ptpmt1-Mast3 (ρ=0.97), Dusp1-Egfr (ρ=0.73), Dusp1-Mapk8 (ρ=0.66), Dusp16-Mapk8 (ρ=0.83), Dusp16-Lrrk2 ρ= (0.86), Dusp10-Igf1R (ρ=0.99).

To identify the interacting partners of DUSPs in TLR4 signaling, we carried out interactome analysis of members of the DUSP and protein kinase families that were found to be differentially expressed in dendritic cells and monocytes (**Supplementary Figures 6**, **7**, **and 8**). Analysis of network properties identified several proteins that seemed to be regulated by DUSP signaling. In the networks of differentially expressed proteins identified in activated mDCs, proteins including Akt1, Lrrk2, Pim1, Dusp1, Dusp16 (overexpressed), Mapk1/3, Pik3cg, Cdk1,Dusp3 and Dusp19 (downregulated) had high number of edges and high betweenness centrality suggesting their importance in innate immunity. In activated hDCs, proteins including MAPK1, CDK2, PIK3CA, MAPK13, DUSP1, DUSP5, DUSP7 and DUSP10 (upregulated), PRKACA, LRRK2, CHUK and DUSP13 (downregulated) had a high betweenness centrality and were identified to be relevant in LPS-induced signaling. Similarly in activated hMOs, PRKCZ, CDK6, DUSP10, TGFBR1 (upregulated), LRRK2, ATM, ATR, MAPK1, DUSP1, MAP3K1 (downregulated) among others had a high betweenness centrality.

## 3. Discussion

Dual-specificity phosphatases (DUSPs) are a family of phosphatases that can act on both serine/threonine and tyrosine residues of several protein substrates leading to wide-ranging effects on cellular signaling and biological processes. With the exact number of DUSP members still being controversial [7, 58, 59], we chose to consider the latest classification described by Chen et al [7], consisting of 40 DUSPs with 9 subfamilies containing more than one member. We validated the subfamily-based classification of DUSPs using two approaches namely-sequence alignment and SMART-based domain analysis. Evolutionary conservation analysis of DUSP family members revealed high sequence conservation of all DUSP members especially in higher mammals.

To determine the extent of expression of dual specificity phosphatases and protein kinases across the human hematopoietic cells and lymphoid organs, publicly available datasets were mined. Since hematopoietic cells are derived from lymphoid tissues, we chose to analyze expression profiles of these. To date, Rieckemann *et al.* have provided the largest expression dataset pertaining to 28 different hematopoietic cell types consisting of high throughput omics data acquired under a single platform containing. Our analysis of this dataset revealed new insights into the expression dynamics of several understudied DUSPs such as slingshot family of phosphatases (SSH1, SSH2 and SSH3) and STYX in various hematopoietic cell types. Additionally, our analysis also resulted in the evaluation of the expression patterns of several understudied protein kinases described by Huang et al. [60] including STK17A, SCYL3, MAST3, CSNK1G2, and the RIO family of kinases (RIOK1, RIOK2 and RIOK3). Furthermore, among the hematopoietic cells, erythrocytes and thrombocytes (platelets) expressed the least number of DUSPs and PKs The restricted expression patterns of these proteins could be potentially exploited for therapeutic modalities in erythrocyte and platelet disorders. In fact, four different PKs (PRKCD, KALRN, MYLK and PTK2) showed restricted expression in thrombocytes. PRKCD has been previously reported to modulate collagen-induced platelet aggregation [61] while MYLK and PTK2 have been reported to be important in megakaryopoiesis [62, 63]. Further, the different patterns of DUSP and PK expression in erythrocytes and thrombocytes compared to the rest of the hematopoietic cells may be attributed to their structures, diverse biological functions performed by these cells, their numbers in the human body and the differing lineages that form erythrocytes and thrombocytes during hematopoiesis.

Among the 701 DUSP-kinase pairs with potential reciprocal activities identified through our analysis, several pairs with previously known reciprocal actions were also identified thereby proving our analysis methods. Among these, SSH1-AURKA pair was identified with a high Spearman’s rank correlation coefficient (ρ= 0.94). Aurora Kinase A (AURKA) overexpression has been previously reported to increase the expression of slingshot kinase 1 (SSH1) resulting in increased cofilin activation and migration of breast cancer cells [64]. Other notable pairs that we identified from the correlation and that were previously reported in the literature included DUSP1-MAPK7 (ρ= 0.79) and DUSP1-MAPK8 (ρ= 0.82). DUSP1 gene silencing has been shown to increase the expression of MAPK7 and MAPK8 transcripts in osteosarcoma cells suggesting reciprocal actions between them [65]. DUSP10 (MKP-5) has been well known to dephosphorylate MAPK8 (JNK) [66] and has also been implicated in immune function. Knocking down DUSP10 expression increased JNK activity and inflammation in murine mesangial cells while its overexpression led to decreased JNK activation [67]. A study by Zhang *et al*. further found that Dusp10-deficient murine cells exhibited increased JNK (MAPK8) activity, elevated levels of proinflammatory cytokines and increased T cell activation [68]. In our correlation analysis, the DUSP10-MAPK8 pair was identified with a Spearman’s rank correlation coefficient of 0.81

We used a subset of omics datasets to investigate the expression of DUSPs and their interplay with PKs upon LPS stimulation in activated dendritic cells and monocytes. Though a previous study on the meta-analysis of TLR4 signaling datasets exists, its focus was on activated macrophages [69]. We also correlated DUSP and PK expression in these cells and carried out interactome analysis to identify key molecules that are potential regulators of LPS-induced signaling. Correlation analysis indicated the presence of DUSP-PK pairs with reciprocal activities in response to activation. Some of these pairs including DUSP1-MAPK8, DUSP1-MAPK8 have already been described in previous literature, thus confirming our findings. DUSP-16 (MKP-7) has previously been identified to regulate MAPK8 (JNK) in LPS-activated macrophages [70] and in activated endothelial cells [71]. Dusp16 (MKP-7) was also reported to have a critical role in the activation and functioning of T cells. Dusp16-deficient T cells had an exaggerated response to TCR activation and had enhanced proliferation properties.[72] DUSP16-deficient macrophages have also been reported to overproduce IL-12 in the context of TLR stimulation [73]. DUSP1 has been reported to play a key role in the feedback control and regulate MAPK8 during glucocorticoid-mediated repression of inflammatory gene expression [74]. DUSP1 is already known to regulate the expression of LPS-induced genes [14]. We also found DUSP 1 and DUSP16 to be important regulators of several MAP kinases from the interactome analysis.

Several novel DUSP-PK pairs were identified and these require further characterization to confirm their role in immunity. Particularly interesting among these include pairs involving slingshot phosphatases which constitute a group of understudied phosphatases. SSH1-CDK13, SSH1-CDK19, SSH2-EPHB2 were identified with high significance. Slingshot phosphatases have been previously implicated in cancer progression [75, 76] and have been known to mediate caspase-modulated actin polymerization towards bacterial clearance upon *Legionella* infection [77]. In our analysis, Slingshot phosphatase members SSH1, SSH2 were found to be downregulated exclusively in human DCs, while SSH3 was downregulated in both human DCs and MOs. Slingshot phosphatase members SSH1 and SSH2 were found to be downregulated exclusively in human DCs, while SSH3 was downregulated in both human DCs and MOs. DUSP10 and IGF1R expression were also found to be highly correlated. Insulin-like growth factors have been known to inhibit anti-tumoral responses of dendritic cells through the regulation of MAP kinases and have therefore been suggested to be targets for immunotherapy [78]. IGF1 has also been reported to influence activation of macrophages in response to high-fat diet or helminthic infection [79]. Additionally, we identified LRRK2 (leucine-rich repeat kinase 2) from the interactome analysis in activated DCs and MOs, to be potentially important in the context of immune signaling. LRRK2 was found to interact with DUSP1 and DUSP16. LRRK2 has been mainly associated with familial cases of Parkinson’s disease [80] and has been known to play an important in innate immunity in the peripheral and central nervous system [81], especially in microglial inflammatory processes [82]. This current finding suggests the possibility of LRRK2 to be additionally important in DUSP-mediated immune signaling.

## 4. Materials and Methods

### 4.1 Datasets

Studies pertaining to immune cells acquired using high-throughput techniques were searched using PubMed. The data matrices pertaining to each dataset were downloaded from the site of the publisher of these articles. Gene expression datasets were downloaded from Gene Expression Omnibus (GEO, https://www.ncbi.nlm.nih.gov/geo/) wherever applicable. The details of all the studies used in this study are provided in **Supplementary Table 1.**

### 4.2 DUSP and kinase lists for analysis

The list of DUSPs used for the analysis was sourced from Chen *et al* [7]. The list of human and mouse kinases was sourced from UniProt (https://www.uniprot.org/docs/pkinfam) which used data from [4, 83, 84]. The master list of dual specificity phosphatases and PKs used for data analysis in the current study are provided as **Supplementary Tables 2 and 3**.

### 4.3 Classification of DUSP family members, **domain analysis and species conservation analysis of DUSP sequences**

The similarity tree between DUSPs was drawn using iTOL with alignment performed with Clustal Omega. Briefly, RefSeq accessions of the longest protein isoforms for all dual specificity phosphatases were retrieved from NCBI gene (https://www.ncbi.nlm.nih.gov/gene). The protein sequences were obtained with Batch Entrez (https://www.ncbi.nlm.nih.gov/sites/batchentrez) and aligned using Clustal Omega (https://www.ebi.ac.uk/Tools/msa/clustalo/) using default settings. The output alignment in PHYLIP (.ph) format and was visualized with Interactive Tree of Life (https://itol.embl.de/) with custom colors and tracks.

Domain analysis was carried out for the longest isoforms of all proteins belonging to DUSP subfamilies with multiple members using the SMART domain prediction tool (http://smart.embl-heidelberg.de/) [85], using the option “PFAM domains”. Orthology data for all human genes were obtained from Homologene (Release 68, downloaded on October 4, 2018 from https://www.ncbi.nlm.nih.gov/homologene) for the analysis of sequence conservation across species. The counts for all genes in the Homologene database were obtained and the Taxonomy ID for each gene was mapped to the species type. The densities of ortholog counts for DUSP family members was plotted against the density of ortholog counts for all human genes in the background using R (v3.5.1).

### 4.4 Landscape of DUSPs and kinases in immune cells

Proteomic and transcriptomic data matrices were obtained from supplementary files of respective articles and accessions were converted into Entrez gene accession formats using bioDBnet:db2db (https://biodbnet-abcc.ncifcrf.gov/db/db2db.php) [86] and g:Profiler (https://biit.cs.ut.ee/gprofiler/gconvert.cgi) [87]. Gene expression data was also downloaded from Gene Expression Omnibus (GEO) wherever supplementary data was not available. Z-score-based normalization of data matrices from all studies was carried out using base R (v3.5.1). DUSP and kinase expression data were subsequently obtained from the normalized datasets. Cell type data and cell sorting information wherever available were retrieved for each of the expression datasets and appended with the expression data. Heatmaps were drawn in Morpheus (https://software.broadinstitute.org/morpheus/) with hierarchical clustering based on Euclidean distance metric, complete linkage method and clustering by rows and columns. We carried out correlation and interactome analysis to look at the interplay between DUSPs and PKs in naïve and activated immune cells. Correlation analysis between z-scores of kinase and DUSP profiles of immune cell proteomes was performed using “Spearman” method through R (v3.5.1). Heatmaps were drawn using Morpheus (https://software.broadinstitute.org/morpheus/).

### 4.5 Landscape of DUSPs and kinases in primary and secondary lymphoid organs

We compiled tissue-based expression data from various datasets present in the Expression Atlas (https://www.ebi.ac.uk/gxa/home). Briefly, the pre-processed datasets from The FANTOM5 project [47], Genotype-Tissue Expression (GTEx) Project [48], The Human Protein Atlas [49, 50], Illumina Body Map [51], NIH Roadmap Epigenomics Mapping Consortium [52] and the ENCODE project [53] were downloaded (Supplementary Table 1, Download date January 11,2019). The datasets were subjected to z-score-based normalization using R (v3.5.1). Data from primary and secondary lymphoid tissues were selected and DUSP and kinase expression data were compiled using in-house scripts. The data was plotted as heatmaps with Morpheus using the same parameters as in the previous section.

### 4.6 Baseline DUSP Interactome

We analyzed publicly available Protein-protein interaction (PPI) data to identify DUSP-kinase interactions. We chose the comPPI database (Compartmentalized Protein-Protein Interaction Database, v2.1.1, http://comppi.linkgroup.hu/home) to identify biologically significant high-confident interactions between proteins with similar subcellular localization patterns. ComPPI is an integrated database of protein subcellular localization and protein-protein interactions from multiple databases including BioGRID, CCSB, DIP, HPRD, IntAct and MatrixDB. [88]. The highly confident interactomes of each member of the dual specificity phosphatase family for *Homo sapiens* were fetched from comPPI, filtering for localization score and interaction score thresholds of 0.7 each. The accessions of interacting proteins were obtained through bioDBnet: db2db (https://biodbnet-abcc.ncifcrf.gov/db/db2db.php) and g: Profiler (https://biit.cs.ut.ee/gprofiler/gconvert.cgi) and the set of interactions were compiled and visualized in Cytoscape (version 3.7.0) to obtain an integrated DUSP interactome. The interactome was clustered into DUSP neighborhood networks using the AutoAnnotate (v1.2) package. The network statistics of the interactome were analyzed using the Network Analyzer tool in Cytoscape [89]. The betweenness centrality measure was used to identify protein hubs central to the DUSP network, which could influence the flow of information triggered by DUSPs. Individual interactomes of each DUSP member were also analyzed to identify central proteins associated with DUSPs using the betweenness centrality measure.

### 4.7 Landscape of DUSPs and kinases in activated immune cells

Data matrices from proteomics and transcriptomics datasets containing gene/protein expression data were fetched from the supplementary tables or GEO for the following studies [29, 32, 33, 55]. DUSP and kinase expression data were obtained from the normalized datasets. Fold-change ratios were calculated by dividing the ratio of intensity/RPKMs of activated/stimulated cells by the ratio of respective intensity/RPKM values of steady-state/unstimulated cells. The fold-change ratios were converted into log2(fold-change) by log transformation. Genes/proteins with log2(fold-change) of 1 (fold-ratio of 2) were considered to upregulated while those with -1 (fold-ratio of -1) were considered to be downregulated. Genes/proteins with ambiguous trends across studies were ignored (both up and down) and only those with overall trends of either up and down were considered for further analysis and genes. We carried out correlation and interactome analysis to investigate the interplay between DUSPs and PKs in activated immune cells. Correlation analysis between z-scores of kinase and DUSP expression profiles of activated immune cells was performed as described above.

### 4.8 Functional, **pathway and network analysis**

The differentially expressed molecules in response to LPS were analyzed with Gene Ontology-based functional analysis through CLUEGO (ClueGO v2.5.2 + CluePedia v 1.5.2) [90] in Cytoscape (v 3.7.0) to identify processes affected by LPS. The parameters used included ‘ClueGO: Functions’ analysis mode, murine ‘GO: Biological Process’ (dated 18.01.2019), global network specificity, pV <= 0.05000 with GO Term grouping. Hypergeometric enrichment-based pathway analysis was performed using Reactome Pathways [91] in R/Bioconductor 3.5.1/3.7 [92] with clusterProfiler 3.8.1 [93] and reactome.db 1.64.0. Genesets with less than 10 genes were excluded from the analysis and p.values were adjusted by Benjamini-Hochberg correction. Pathways reaching adjusted p-values <= 0.0075 were curated manually and plotted with ggplot2 3.1.0. Network analysis of activated DCs and MOs was performed using STRING in Cytoscape. The network properties were calculated using Network Analyzer.

## 5. Conclusions

Though the role of dual specificity phosphatases in innate and adaptive immunity is known, their interplay with kinases was not precisely understood. In the current study, we expanded the knowledge on the role of dual specificity phosphatase signaling in activated and steady-state cells through the analysis of high-resolution expression datasets. We confirmed the importance of several known DUSPs such as DUSP1 and DUSP10 in innate immunity. We also report potentially novel role of DUSPs such as the Slingshot phosphatases and PKs such as LRKK2 in immune signaling. These need to be further validated to confirm their roles. we also identified selective patterns of expression of a few DUSPs and PKs across hematopoietic cells which could be used as potential therapeutic targets. Furthermore, we also identified potential species-specific events of DUSP signaling which need to be further validated. Finally, we demonstrate the utility of meta-analysis of existing datasets to identify molecular mechanisms of various biological processes and fill existing gaps in understanding understudied proteins. The findings from this study will aid in the understanding of DUSP signaling in the context of innate immunity.

## Author Contributions

Conceptualization, R.K.K.; methodology, R.K.K.,Y.S.; formal analysis, Y.S., K.B.; data curation, S.P.; writing—original draft preparation, Y.S.; writing— review and editing, S.P., KB. R.K.K., T.S.K.P. supervision, R.K.K. and T. S. K.P.; funding acquisition, R.K.K.”,

## Funding

This work was funded by the Research Council of Norway (FRIMEDBIO “Young Research Talent” Grant 263168 to R.K.K.; and Centres of Excellence Funding Scheme Project 223255/F50 to CEMIR), Onsager fellowship from NTNU (to R.K.K.).

## Acknowledgments

We thank the authors of the datasets used in this study for making their data publicly available.

## Conflicts of Interest

The authors declare no conflict of interest

## Abbreviations

DC: Dendritic cells
MO: Monocytes
LPS: Lipopolysaccharides
DUSP: Dual Specificity Phosphatase
PK: Protein Kinase
GO: Gene Ontology

### Appendix A Supplementary Tables

**Supplementary Table S1.** List of datasets used for the study

**Supplementary Table S2.** List of DUSP family genes used for the study and their details

**Supplementary Table S3.** List of protein kinases used for the study and their details

**Supplementary Table S4.** Expression of kinases and DUSPs (z-scores) in hematopoietic cells (matrix from Rieckmann et al, Nat Immunol (2017))

**Supplementary Table S5.** Matrix for corresponding RNA and protein expression data (z-scores) for T4 naïve, T4 TCM, B memory and classical monocyte cells

**Supplementary Table S6.** Expression of DUSPs and protein kinases in primary and secondary lymphoid tissue expression datasets.

**Supplementary Table S7.**Correlation matrix between DUSP and protein kinase expression in hematopoietic cells (data from Rieckmann et al, Nat Immunol (2017)). Spearman’s rank correlation coefficient was used.

**Supplementary Table S8. List of nodes and their properties from the comPPI interactome for DUSP family members.** The interactions were obtained from the comPPI database and the network properties analyzed using Network Analyzer in Cytoscape.

**Supplementary Table S9. Expression of kinases and DUSPs in activated hematopoietic cells (matrix from Rieckmann et al**, **Nat Immunol (2017)).** Ratios obtained by dividing expression values in the activated state by expression values in the steady-state.

**Supplementary Table S10. log2(fold-change) of kinases and DUSPs in LPS-stimulated hematopoietic cells.** Ratios obtained by dividing expression values in the activated state by expression values in the steady-state and log transformation to the base 2. The study code indicated in the header can be referred to in Supplementary Table 1. The kind of biomolecule assayed and the time point after LPS stimulation is mentioned in the header.

**Supplementary Table S11.** List of DUSPs and kinases differentially expressed in monocytes and dendritic cells in response to LPS

**Supplementary Table S12.** List of enriched pathways containing proteins differentially expressed in monocytes and dendritic cells in response to LPS

**Supplementary Table S13.** Correlation matrix between DUSP and protein kinase expression in activated DCs and MOs. Spearman’s rank correlation coefficient was used.

**Supplementary Figures**

**Supplementary Figure 1. Baseline expression of A.DUSPs and B. protein kinases in hematopoietic cell line proteomic data.** The raw hematopoietic cell expression data obtained from Rieckmann *et al, Nat Immunol* (2017) was scaled to obtain Z-scores. Z-scores were plotted as a heatmap using Morpheus and hierarchical clustering was carried out using Euclidean distance, complete linkage by both rows and columns.

**Supplementary Figure 2. DUSP interaction network**. Protein-protein interaction data between DUSP and other proteins were obtained from Compartmentalized Protein-Protein Interaction (comPPI) Database were analyzed in Cytoscape using Network Analyzer to obtain network properties including Betweenness Centrality. Proteins with high Betweenness Centrality indicate primary regulatory proteins associated with DUSPs. Manual clustering of proteins within the vicinity of DUSPs was carried out using AutoAnnotate 1.3 in Cytoscape.

**Supplementary Figure 3. (a) Differential expression of DUSPs in various activated hematopoietic** cells. (b). Differential expression of protein kinases in various activated hematopoietic cells.

**Supplementary Figure 4. Expression of (a) DUSPs and (b) protein kinases in activated hematopoietic cells.** Expression ratios from activated and steady-state hematopoietic cells were calculated from expression data obtained from Rieckmann *et al, Nat Immunol* (2017) and log transformed. Expression values were plotted as a heatmap using Morpheus and hierarchical clustered by Euclidean distance and complete linkage by both rows and columns. **Similarity matrices for (c) DUSP and (d) protein kinase expression in hematopoietic cells.** DUSP and protein kinase expression in various cell types was correlated via Spearman’s rank correlation.

**Supplementary Figure 5:** Venn diagrams showing the overlap of molecules differentially expressed in response to LPS in **(a).** murine and human dendritic cells (DCs). **(b)**. human dendritic cells and monocytes (upregulated molecules). **(c)** human dendritic cells and monocytes (downregulated molecules). **(d) Correlation of protein kinase and DUSP expression patterns in activated DCs and MOs**. The correlation was carried out using Spearman’s rank correlation method to identify kinase-DUSP pairs that may have reciprocal activities.

**Supplementary Figure 6.** Network analysis of DUSPs and kinases that were (a). upregulated and (b). downregulated in activated murine dendritic cells

**Supplementary Figure 7:** Network analysis of DUSPs and kinases that were (a). upregulated and (b) downregulated in activated human dendritic cells

**Supplementary Figure 8:** Network analysis of DUSPs and kinases that were (a). upregulated and (b) downregulated in activated human monocytes

