## Supplementary material for "Dynamics of dual specificity phosphatases and their interplay with protein kinases in immune signaling": Supplementary_figures.pdf

**Supplementary Figure 8:** Network analysis of DUSPs and kinases that were (a). upregulated and (b) downregulated in activated human monocytes

**Supplementary figure 1**

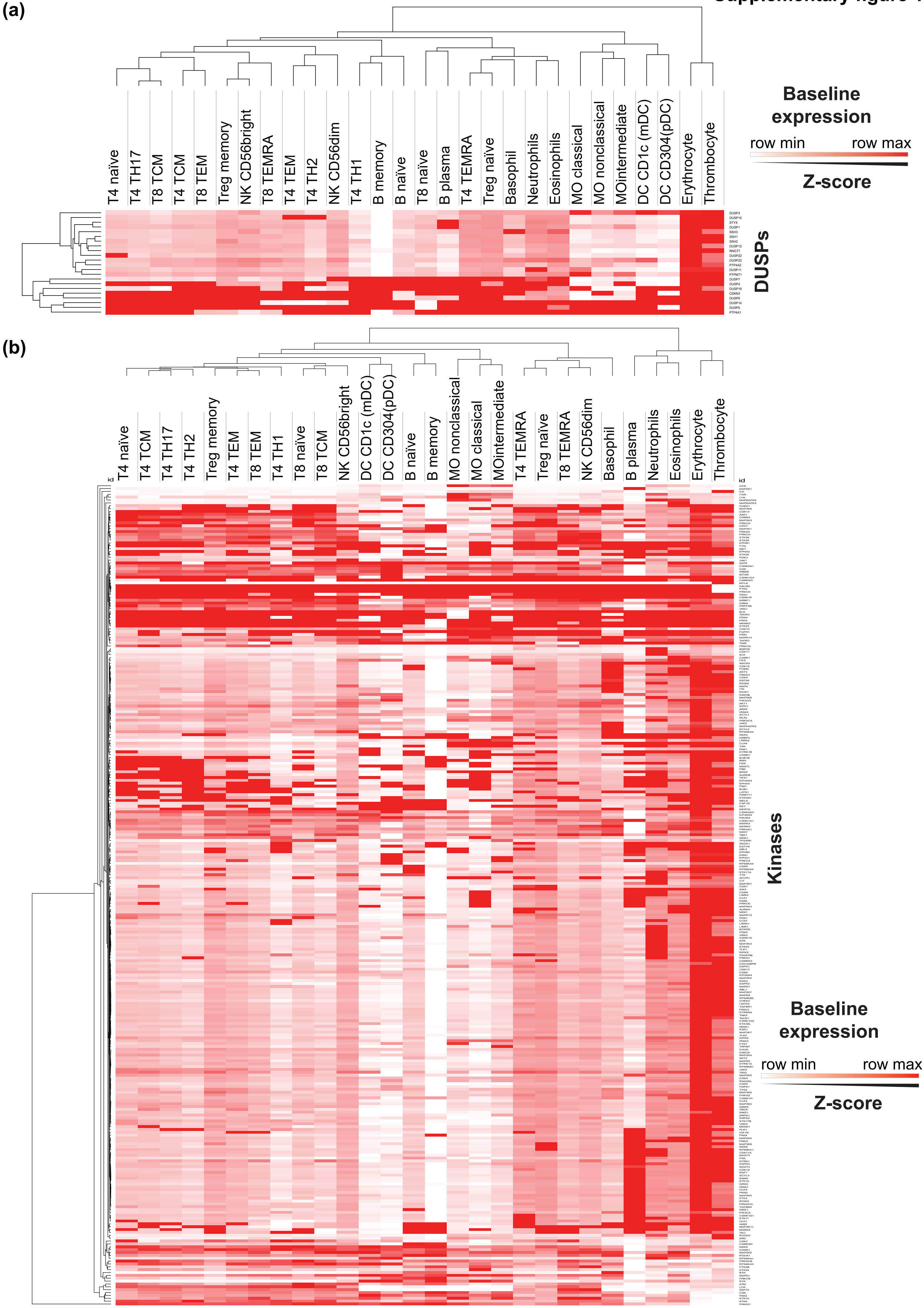

**Supplementary figure 2**

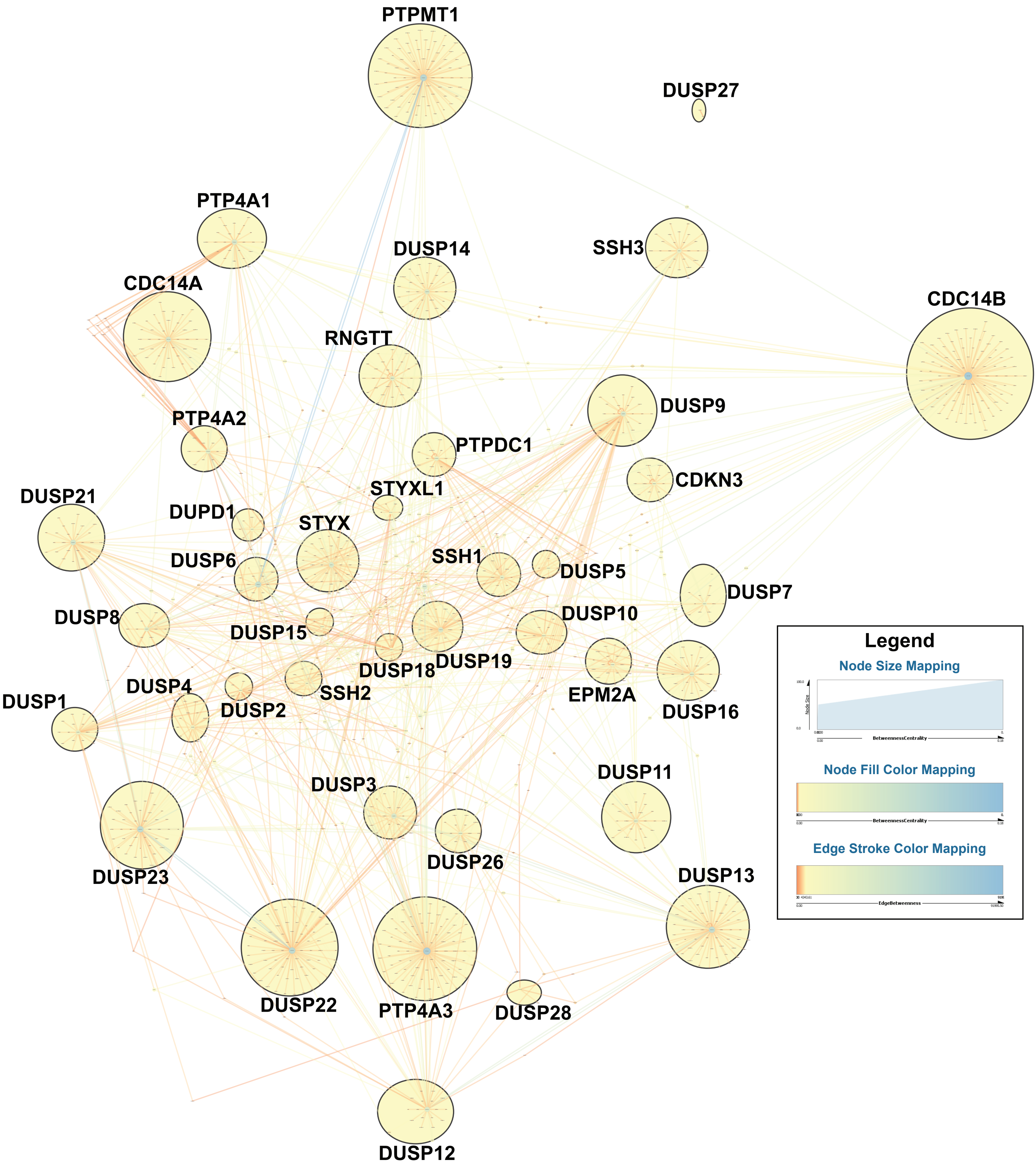

(a)

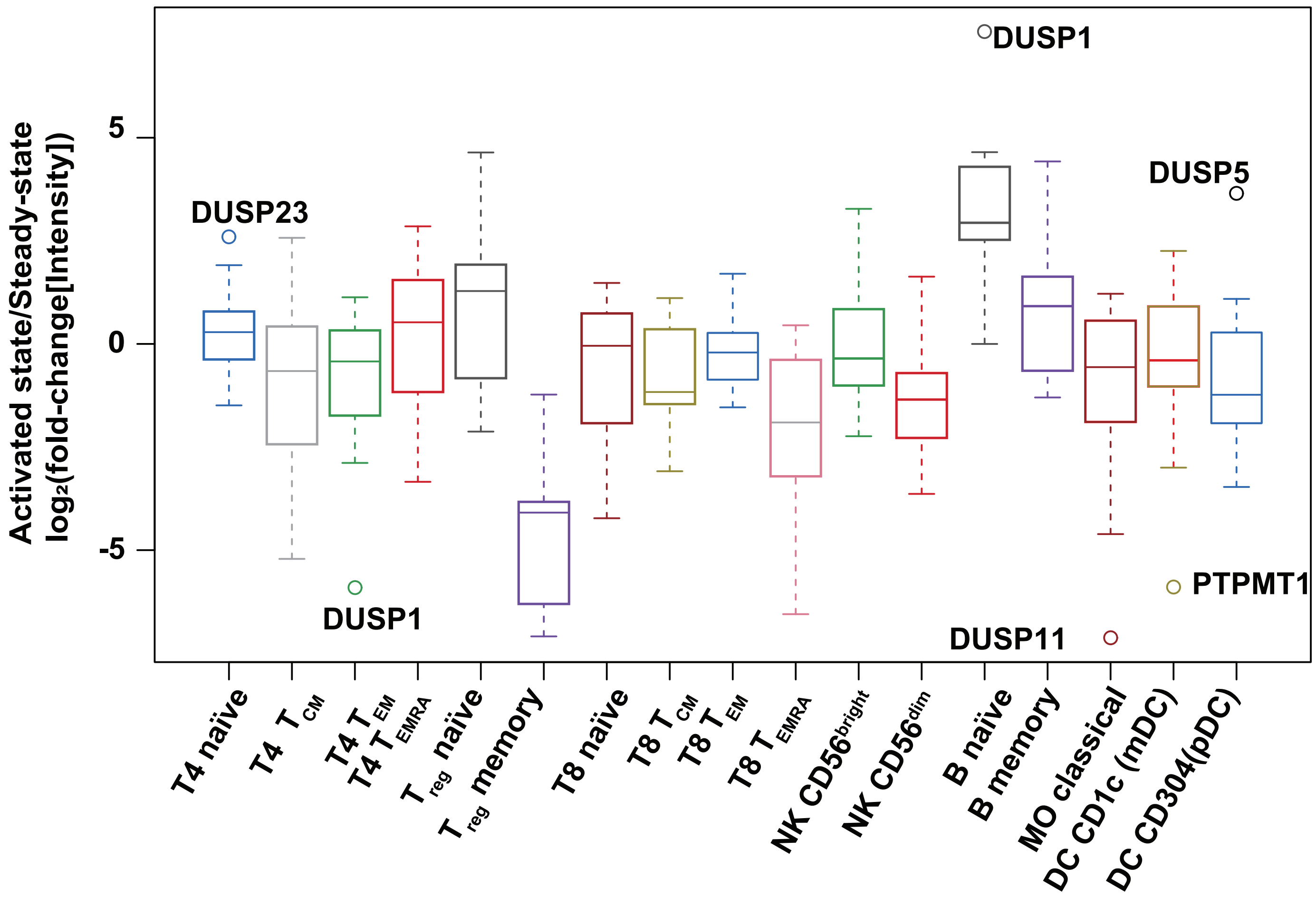
Activated state/Steady-state
log₂(fold-change[Intensity])
5
0
-5
DUSP23
DUSP1
DUSP1
DUSP5
PTPMT1
DUSP11
T4 naïve
T4 T CM
T4 T EM
T4 T EMRA
T reg naïve
T reg memory
T8 naïve
T8 T CM
T8 T EM
T8 T EMRA
NK CD56bright
NK CD56dim
B naïve
B memory
MO classical
DC CD1c (mDC)
DC CD304(pDC)

(b)

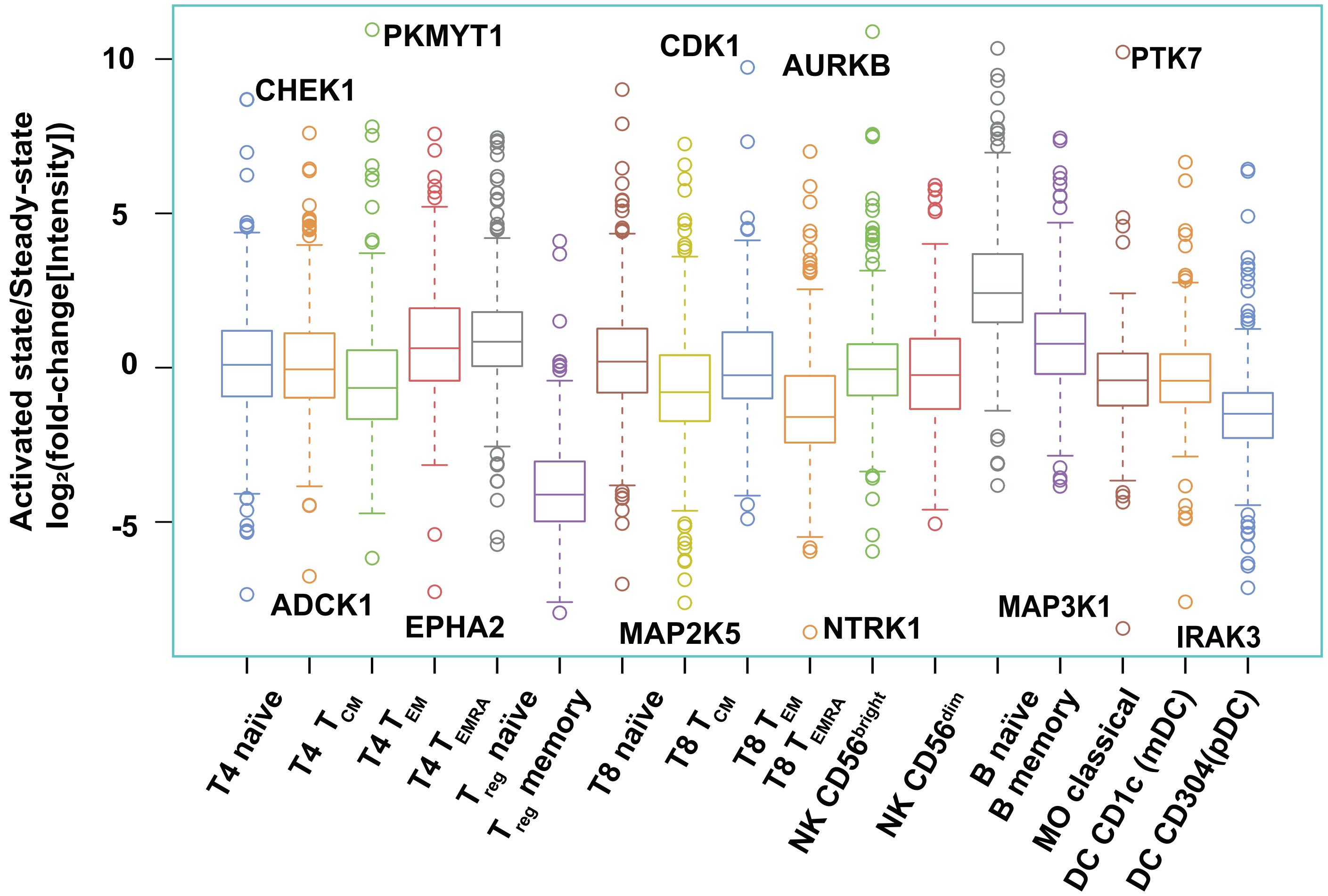
Activated state/Steady-state
log₂(fold-change[Intensity])
10
5
0
-5
PKMYT1
CHEK1
CDK1
AURKB
PTK7
ADCK1
EPHA2
MAP2K5
NTRK1
MAP3K1
IRAK3
T4 naïve
T4 T CM
T4 T EM
T4 T EMRA
T reg naïve
T reg memory
T8 naïve
T8 T CM
T8 T EM
T8 T EMRA
NK CD56bright
NK CD56dim
B naïve
B memory
MO classical
DC CD1c (mDC)
DC CD304(pDC)

Supplementary figure 4

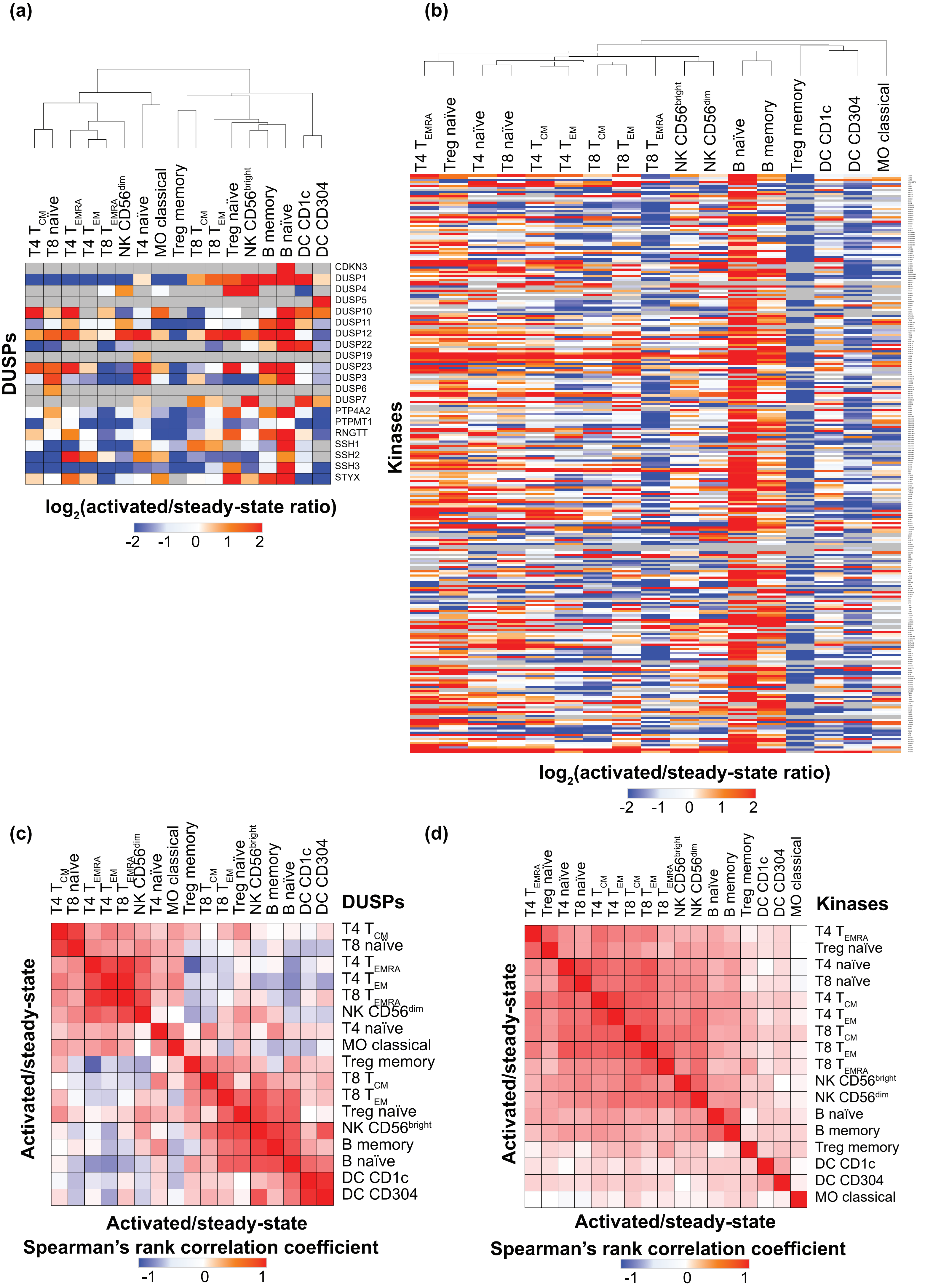

**Supplementary figure 5**

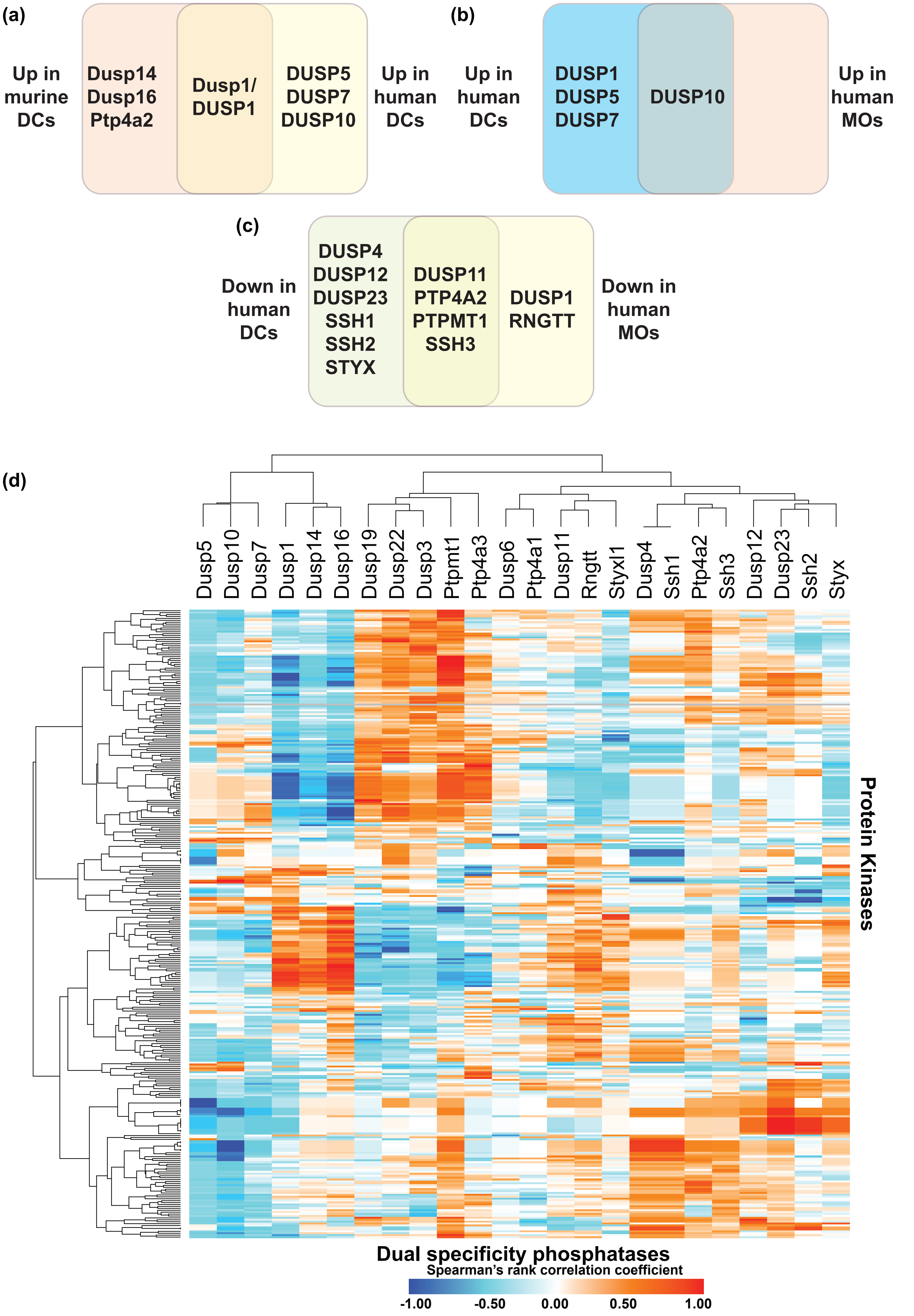

**Supplementary figure 6**

**(a)**

### Upregulated molecules in murine dendritic cells

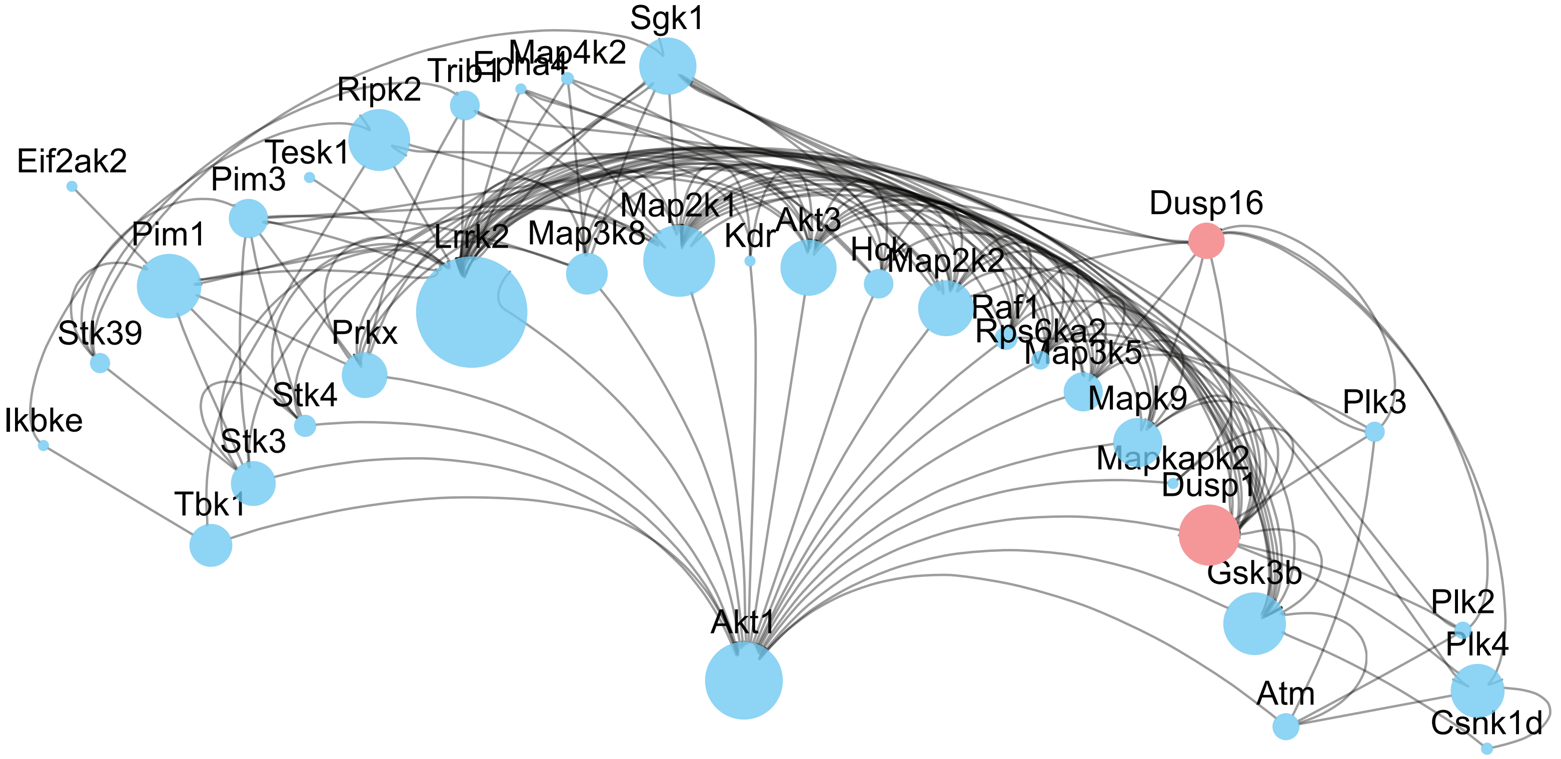


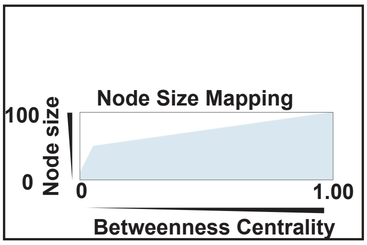


**(b)**

### Downregulated molecules in murine dendritic cells

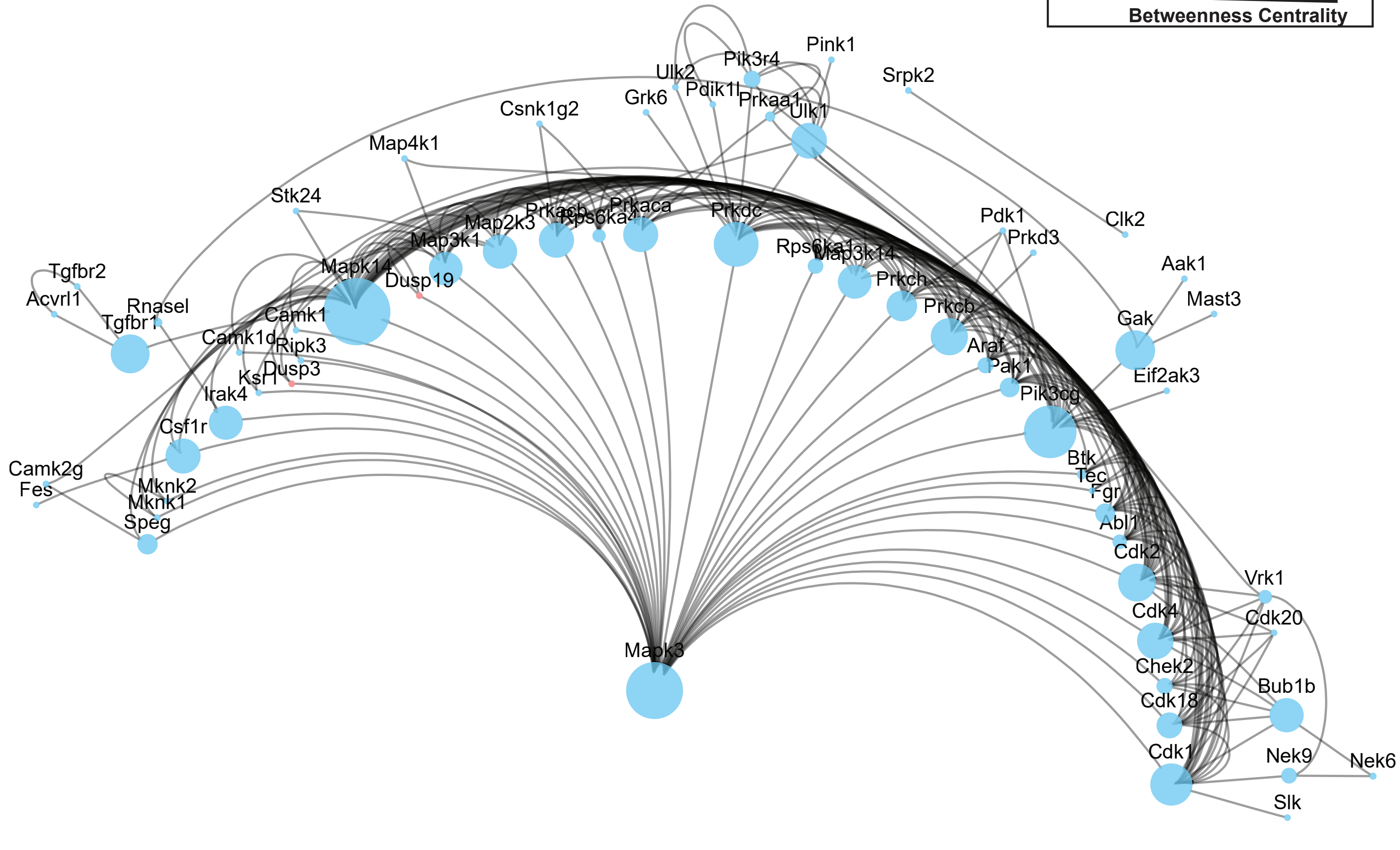

**Supplementary figure 7**

**(a)**

### Upregulated molecules in human dendritic cells

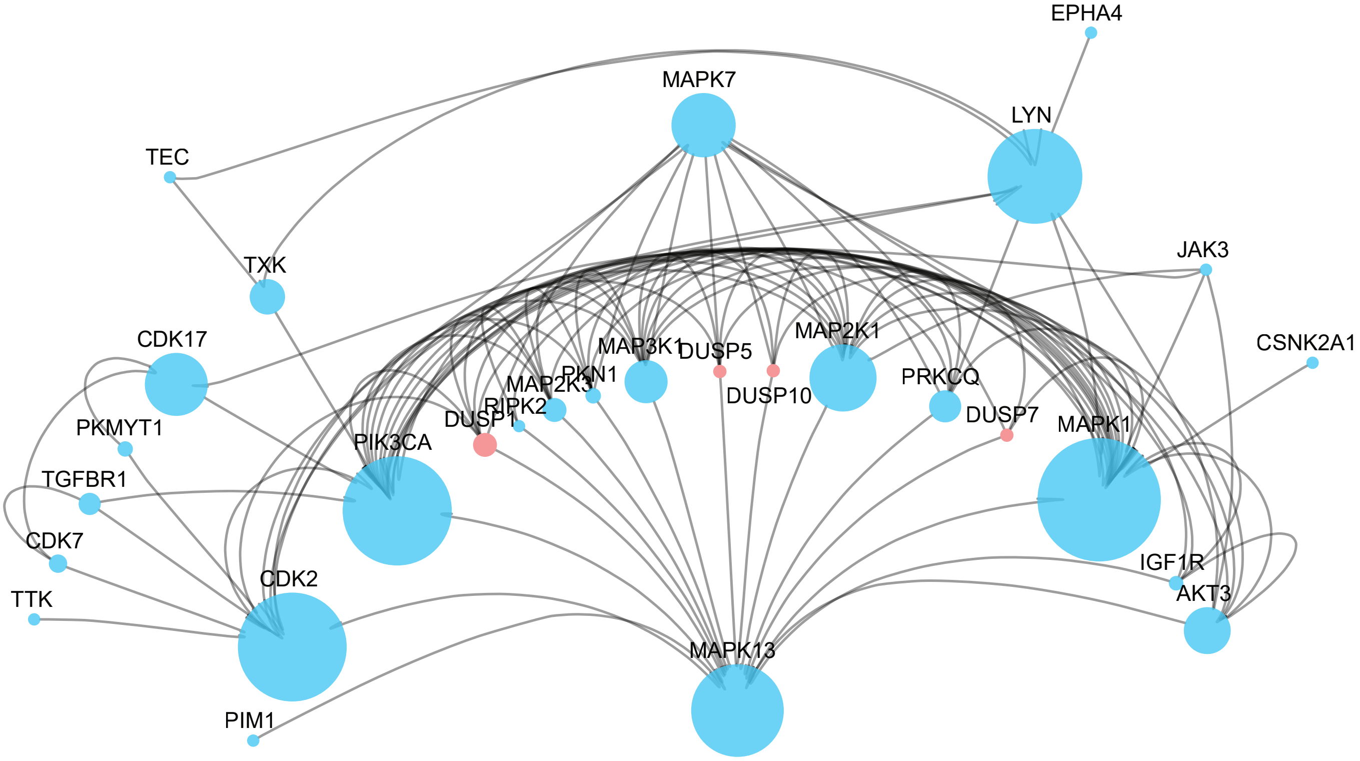


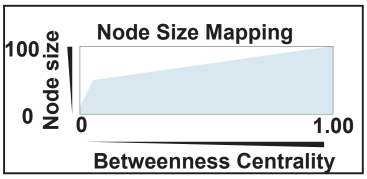


**(b)**

### Downregulated molecules in human dendritic cells

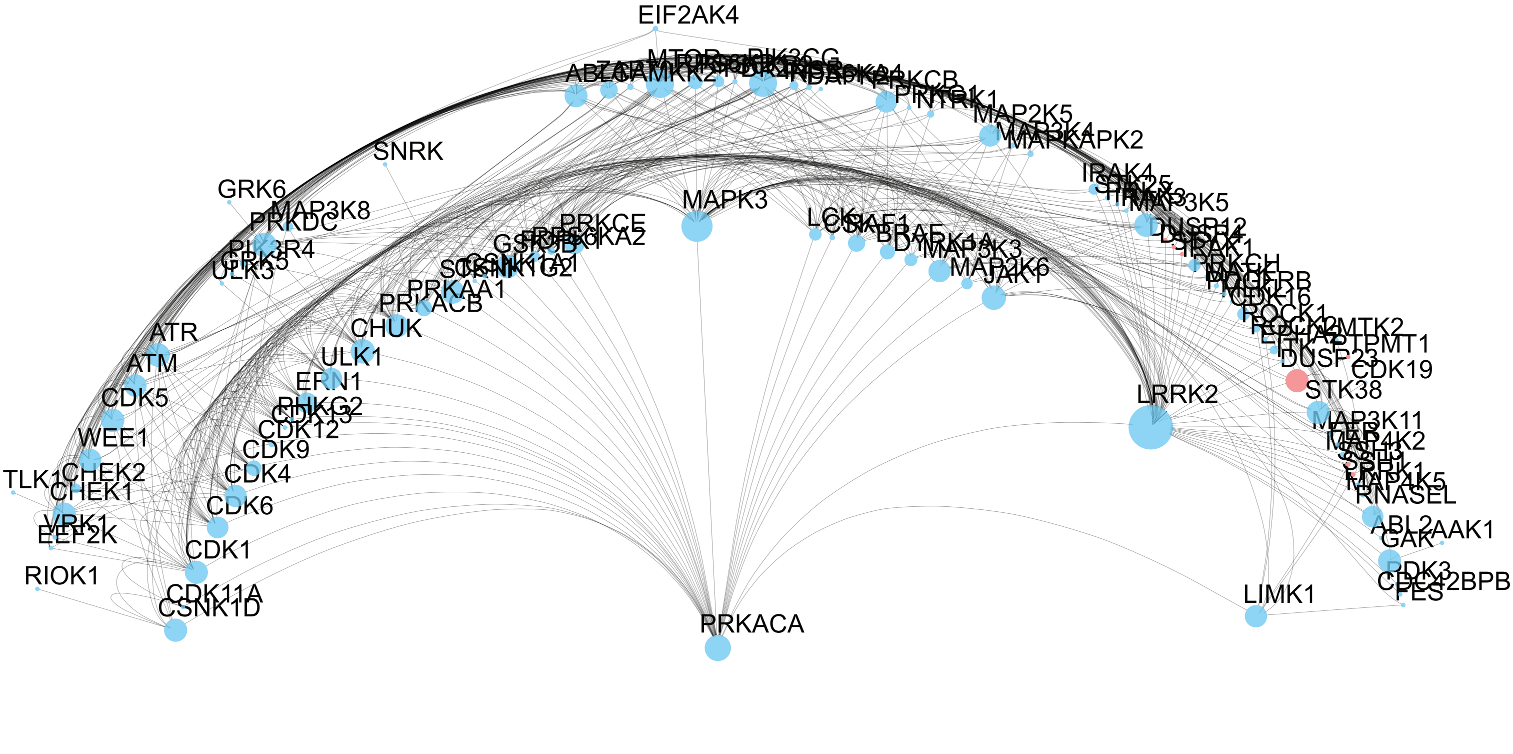

**(a)**

### Upregulated molecules in human monocytes

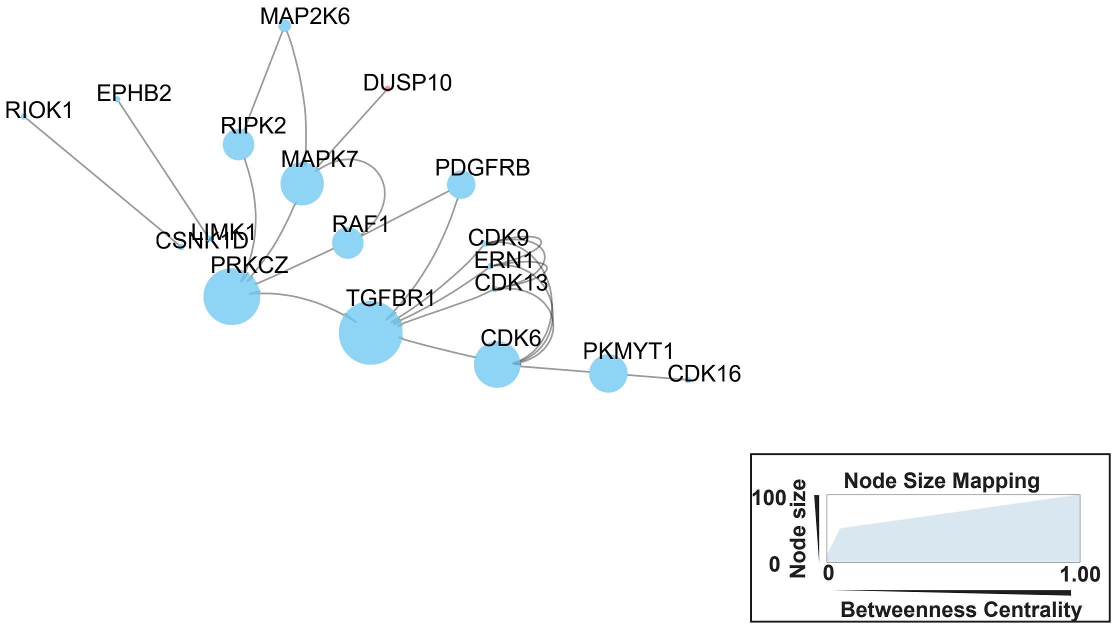


**(b)**

### Downregulated molecules in human monocytes

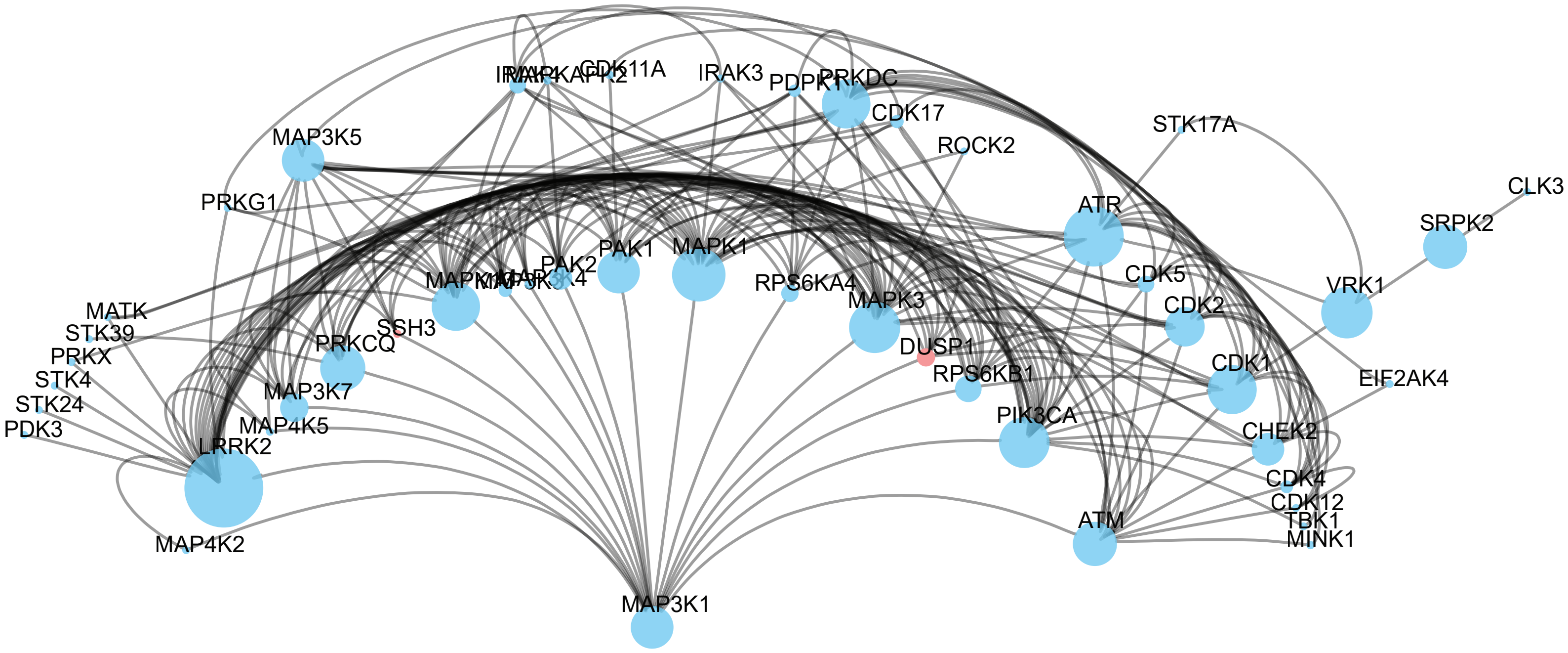
